# The “sewing machine” for minimally invasive neural recording

**DOI:** 10.1101/578542

**Authors:** Timothy L Hanson, Camilo A Diaz-Botia, Viktor Kharazia, Michel M Maharbiz, Philip N Sabes

## Abstract

We present a system for scalable and customizable recording and stimulation of neural activity. In large animals and humans, the current benchmark for high spatial and temporal resolution neural interfaces are fixed arrays of wire or silicon electrodes inserted into the parenchyma of the brain. However, probes that are large and stiff enough to penetrate the brain have been shown to cause acute and chronic damage and inflammation, which limits their longevity, stability, and yield. One approach to this problem is to separate the requirements of the insertion device, which should to be as stiff as possible, with the implanted device, which should be as small and flexible as possible. Here, we demonstrate the feasibility and scalability of this approach with a system incorporating fine and flexible thin-film polymer probes, a fine and stiff insertion needle, and a robotic insertion machine. Together the system permits rapid and precise implantation of probes, each individually targeted to avoid observable vasculature and to attain diverse anatomical targets. As an initial demonstration of this system, we implanted arrays of electrodes in rat somatosensory cortex, recorded extracellular action potentials from them, and obtained histological images of the tissue response. This approach points the way toward a new generation of scaleable, stable, and safe neural interfaces, both for the basic scientific study of brain function and for clinical applications.

## 1. Introduction

Our ability to understand, diagnose, treat, and interact with the brain is in many respects limited by our ability to record and stimulate individual neurons. In addition to optical approaches, surgically implanted stiff metal or silicon electrodes, arranged in rigid arrays, are a standard technology used for this purpose. These work admirably well, yet leave at least four areas open for improvement: *(1)* minimizing the mechanical stress and impedance mismatch between tissue and electrode; *(2)* minimizing overall implant size; *(3)* minimizing vasculature disruption; and *(4)* maximizing the number and anatomical distribution of targeted electrodes. Here we address these four areas by using a single fine and stiff needle to insert many fine and flexible polymer electrodes, each to individually specified targets. The method allows us to obtain maximal device stiffness when penetrating into the brain while retaining maximum flexibility and minimum size of the indwelling implant.

### 1.1. Motivation

Improvements in areas *1-3* serve to collectively reduce the foreign body response (FBR); numerous studies have demonstrated that an electrode elicits a strong FBR when implanted in the brain [1],[2],[3],[4]. The foreign body response causes growth of astroglial and fibrous scar tissue, which ultimately insulates the electrode and pushes neurons outside the recording volume, or outright kills them [5], altogether leading to the electrodes failing. This FBR seems to be initiated by chemical (biocompatibility) and mechanical (micromotion and interfacial stress) [6],[7],[8] means; to remedy it, the material must be biocompatible and subject the brain to minimal mechanical stress.

#### 1.1.1. Mechanical stress and impedance mismatch

There has been much work on minimizing mechanical stress; approaches include not fixing the electrode to the skull or dura [9],[8],[10],[11],[12],[7]; distributing stress along a long track of fine electrodes in “inside-out” arrays [13],[14]; distributing stress between electrodes in an array [15]; making the electrode flexible and inserting with a dissolving stiffener [16][17],[18],[1],[19],[20]; making the electrode dramatically soften upon implantation by coating in an elastic alginate buffer [21] or polymeric nanocomposite [4],[22]; making the electrode surface soft and porous [23]; increasing surface area [3]; or by adding sinusoidal meanders into the electrode [24],[25]. Each of these advances support the hypothesis that minimizing electrode-tissue mechanical stress also minimizes FBR, and hence should maximize electrode performance.

#### 1.1.2. *Size* Mechanical size is also important

beyond displacing less tissue, small implants have a smaller volume (inertia) to surface area (friction) ratio, and thus exert less force upon the brain during accelerations [26]. Likewise, the volume of FBR was shown to be proportional to device cross-sectional area [27]; polymer fibers smaller than 6 *µm* show almost no FBR [28]; 12.5 *µm Ni* − *Cr* − *Al* wires have excellent longevity in monkeys [13],[14]; 800 nm × 20 *µm* SU-8 polymer probes offer many months of recording stability in mice [29]; *<* 10 *µm* carbon fiber electrodes elicit reduced chronic inflammation [30]; carbon fiber arrays yield excellent chronic recording capability [31]. Finally, density differences between probes and tissue, and the resulting inertial forces, seem to cause glial scarring [32]. This motivates us to make the electrodes as thin, flexible, and light as possible and to minimize the coupling between each electrode and dura / cranium while still allowing the electrode to be reliably implanted. Our probes have a density of 1.67 *g/cm*^3^, an order of magnitude lower than tungsten, platinum, or iridium, and 5 times lower than stainless steel, and are 20,000 times more flexible than an equivalent commonly-used 35 *µm* stainless steel microwire or 35*x*50 *µm* silicon shank.

#### 1.1.3. Vasculature

A second reason for recording-site failure is blood-brain barrier (BBB) compromise. The capillary bed in primate cortex is very dense, with microcapilaries spaced at 40 *µm* [33], and nearly a meter of vasculature per microliter [34], hence it is challenging to blindly implant anything without severing or collapsing blood vessels. While surface vasculature is redundant, and can tolerate point infarctions, descending arterioles are much more sensitive to disruption, and hence must be avoided [35]. Meanwhile, other subsurface microvasculature is more robust [36]. Ischemic damage due to vessel blockage can extend several hundred micrometers beyond an electrode shaft [37], results in hemorrhagic necrosis and edema [38], and this leads to highly variable responses, depending on the particular locations of vasculature along electrode tracts [39]. This is a prevalent problem – with a fixed silicon Utah array, 60% of needle tracts showed evidence of hemorrhage and 25% showed edema after a day of implantation [40]; with chronically implanted tungsten microwire arrays, high ferritin expression, indiciative of intraparenchymal bleeding, was found around a subset of all electrodes [41]. Indeed, the importance of BBB compromise with neural interfaces leads some to suggest that the glial scar is an adaptive response to maintain the BBB, and the ED1-reactive cells found around electrodes are partially circulatory monocytes, not microglia [2]. Other studies do not support the astrocyte-encapsulation hypothesis, but rather show that implantation is associated with prolonged BBB permeability, CD68 immunoreactivity, and demylenation [42], and this correlates with animal to animal variability [43]. Either way, the weight of this evidence leads us to try to maximally avoid vasculature while implanting – hence, targeting one at a time under optical guidance.

#### 1.1.4. Targeting

Targeting electrodes requires micron scale accuracy to avoid blood vessels, while retaining the ability to select insertion sites across the milimeter scale surface of the mammalian brain. For example, it would be advantageous to simultaneously record from neurons in different layers of the motor cortex, structures in the basal ganglia, and thalamus; to date, recording such structures simultaneously remains technically challenging [44]. In terms of neuroprosthetics, visual prostheses would benefit from more stimulation sites in the cortex: the user would simply be able to see more clearly. For recording, motor prosthesis would benefit from more electrodes, as there is consistent evidence that averaging over more neurons reduces decoder noise [45],[46],[47],[48],[49]; additionally, more end-effectors or degrees of freedom requires concomitantly more neurons for accurate control. Unfortunately implanting thousands of individually and precisely targeted electrodes, as motivated above, is difficult without automation.

### 1.2. Comparison to current state-of-the art

The sewing machine system solves a number of important practical issues: *(1)* Robotic automation of the insertion process allows each electrode to be efficiently targeted, while also avoiding surface vaculature. Much of the complexity of handling and manipulating micron-scale polymer electrodes is pushed to automation and closed-loop imaging & computer control (Supplemental section 8.1). *(2)* Wiring is managed before insertion (through a controlled release substrate and sequential peel-off) and after (via wicking silicone). Thus, the implant *is* the cabling, minimizing tethering forces, and permitting the backend to be distant from the insertion site. *(3)* Bonding is performed preoperatively, allowing the electrodes to be plated and tested beforehand. In this study, ultrasonic wirebonds are made to a passive adapter, but can easily be extended to active electronics. This has the obvious advantage of reducing surgical time / complexity. *(4)* Both the inserter needle and electrodes reside in cartridges, facilitating preoperative sterilization and intraoperative replacement. Integrated with the robot, this permits high channel count and anatomical coverage.

There are a number of other notable approaches for minimally-invasive neural recording. When compared to syringe-injectable SU-8 mesh electronics [50],[51], our method is less invasive – rather than using a 650 *µm* OD glass capillary to inject, we use a 25 *µm* needle for insertion. Assuming ∼ 600 *µm*^2^ cross-sectional area as the upper limit of our needle and electrode, this means that one syringe injection is first-order equivalent to 553 needle insertions in terms of tissue disruption. Furthermore, we are able to avoid larger superficial vessels, which is more difficult with a large insertion probe. Finally, electrodes are bonded, assembled, and tested pre-operatively. However, our probes are not as thin and flexible as Fu et. al., and have not shown longevity and stability as they have [29].

A second cutting-edge approach uses microfluidic injection of fine carbon-nanotube fibers [52]. This method can be less invasive in that there is no insertion shuttle; however, our approach should reach parity in terms of recording sites / disrupted volume when finer lithography permits 4 or more recording sites per probe. As with syringe-injectable electronics, there is an advantage here in that bonding is done pre-operatively.

The third approach is similar to ours, albeit with a finer carbon-fiber probe, smaller SU-8 electrodes, and significantly better evidence of longevity and minimal scarring [53]. A primary difference here is in scalability through robotization and fabrication; rather than placing the probes on the surface of the brain and carefully inserting them, our probes are peeled off from a pre-assembled and verified cartridge, which keeps them organized prior and during insertion, and allows for a larger number of insertions within a given time. Massey et. al. have a different approach to carbon fiber electrodes; they use a microfabricated silicon supports with *Al/TiN* coated vias and *Ag* ink interconnect to create high-density ultra-fine arrays [54].

Further excellent results have been presented by Jason Chung et al. [20] using polyimide-platinum probes implanted with a stiffener and dissolving adhesive. This method has the advantage that all can be done using stereotatic equipment, and that recordings are stable and durable.

## 2. Methods

The “sewing machine” system consists of three primary sub-systems: microfabricated electrodes, inserter needle, and inserter robot. To better understand how these components interact, we first provide a broad description of the basic implantation procedure, and then discuss each component in more detail.

### 2.1. Overview of insertion procedure

The key elements of the insertion procedure are shown in Figure 1. For in-vivo studies in rat, stereotaxic technique is used, and a craniotomy is performed to expose the dura mater (further surgical details are given below in section 2.4.1 *Surgical Methods*). An image of the surface of the brain (or agar proxy) is captured with a surface-imaging camera mounted to the surgical robot. Based on this image, individual target sites are chosen via custom software. Targeting is based on the anatomically specified target region, the desired density of insertions, and to avoid surface vasculature. Laser-ablation then creates targeted microdurotomies at each insertion site. Next, an electrode cartridge is loaded into the robot, which positions the cartridge adjacent to the selected target region (in rats, the cartridge is posterior to the craniotomy.) Electrodes are fabricated with a small loop on the end, which allows the inserter head to grasp the tip of an individual electrode by engaging the needle in this loop. The robot then peels electrodes off the cartridge, moves over the brain, and inserts them. The needle is then removed, leaving the electrode in place, and the process repeats. See figure 5 for a depiction of the peel-off and insert steps. Once all electrodes in a given array are inserted, the reference is placed on the surface of the brain, and all loose shanks are protected with silicone gel. Then the PCB to which they are wirebonded is released from the electrode cartridge, and this PCB, to which external connectors are mounted, is positioned over the craniotomy and fixed in place with dental acrylic.

**Figure 1:**
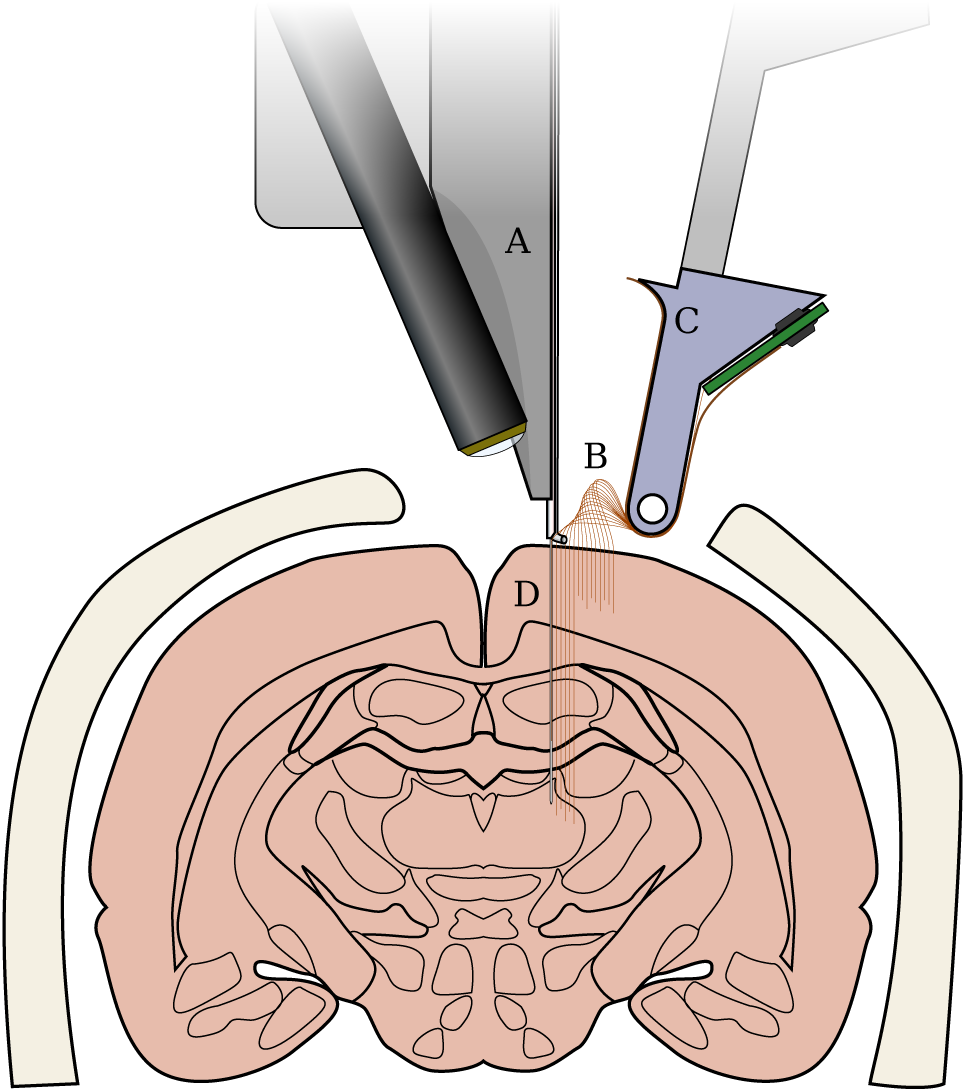
Not-to-scale schematic of the “sewing machine” method for inserting fine and flexible electrodes into the brain. Inserter head, **A**, moves in three dimensions to individually pick flexible electrodes **B** from a replaceable, sterilizable cartridge **C**. Electrodes are then inserted with a fine needle **D** into the brain. The needle is retracted, leaving the electrode in place, and the inserter head moves to pick a new electrode. Electrodes and needle are enlarged to make them visible here.

### 2.2. Electrodes

Arrays consisting of 64 individual electrode threads were lithographically fabricated using polyimide as the substrate and platinum as the conductor. Each thread has a single recording or stimulating site and a loop at the end, which serves as the engagement point for a needle that is used to position and insert the thread. Some electrodes have a pair of ‘dog-ears’ or barbs for holding the electrode in place in the tissue. The elements are all shown in Figure 2 B,D. The recording or stimulating site was wired to bonding pads through a 4*µm* wide trace on a 16*µm* wide shank, as shown in Figure 2 C,D.

**Figure 2:**
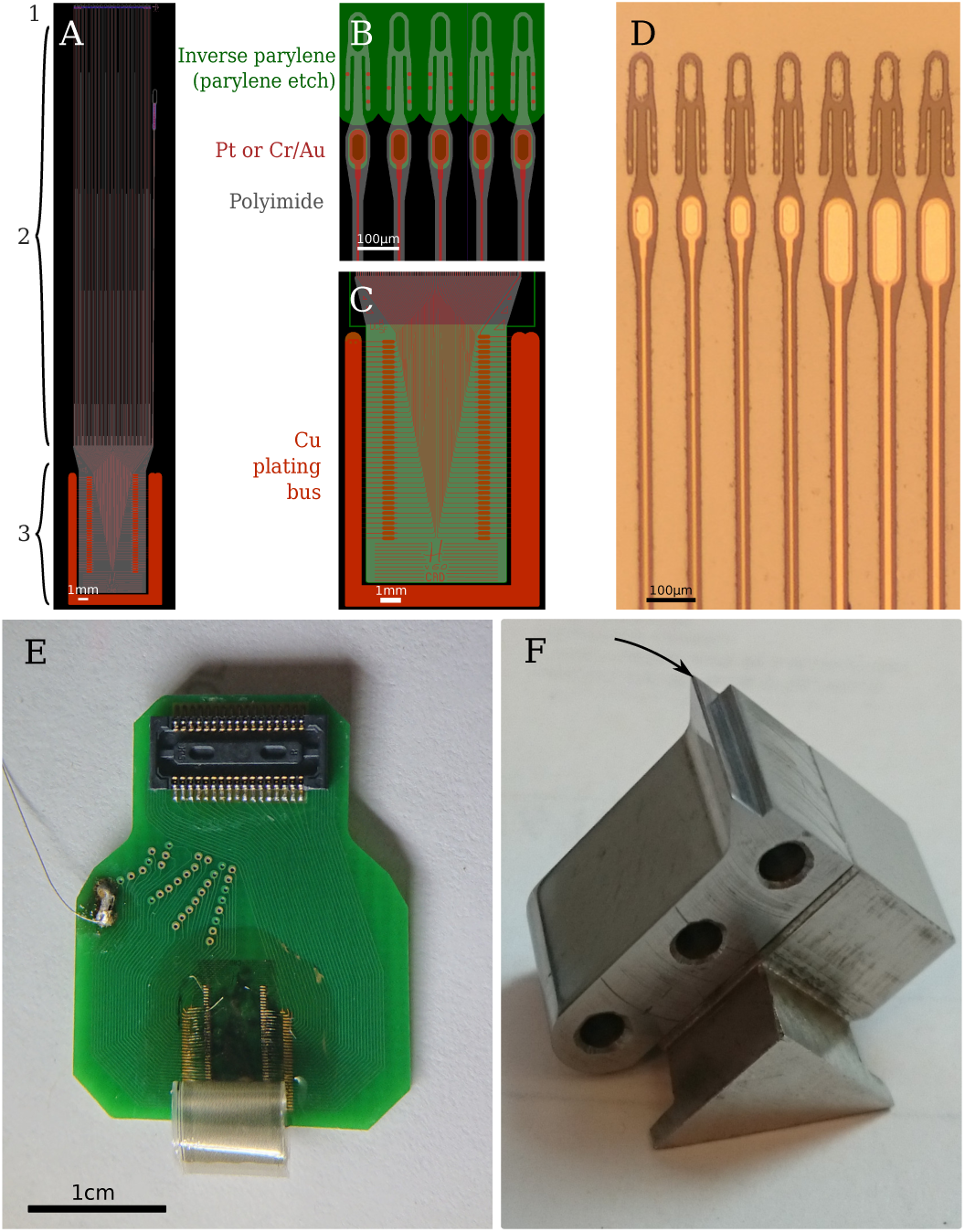
Electrodes and cartridge. **A**. Overview of one electrode array in the design program, Kicadocaml. Red is metal (Pt), gray is device outline (polyimide), green is inverted parylene (parylene etch). *1.* Recording site and loop, detail in B. *2.* Tapered electrode shank. Total length is 27.25 mm. *3.* Bondpad area, detail in C. Bondpad size is 4.2 × 7.7 mm. To the right is the reference electrode, which is peeled off the electrode cartridge last by way of the large loop and fine forceps. **B**. Detail of the electrode head, showing the elongated loops, recording sites, and binary electrode numbering along the dog-ears. Green again is parylene – the loops and protrude above the parylene edge. Note also holes in parylene for electroplating. **C**. Detail of bondpad area. Large area of red-orange is the Pt/Cu bus, which supplies current to the wirebond sites (ovals) through 10 *µm* × 115 *µm* Pt/PI bridges. This bus is peeled away before releasing the electrodes from the wafer. **D**. Micrograph of fabricated electrodes prior parylene deposition. **E**. Photograph of electrodes, epoxied and wirebonded onto a connector PCB. Electrodes are curled due to stress in the parylene. **F**. Photograph of unpopulated electrode cartridge. Knife-edge, where loops are exposed, indicated with arrow.

Polyimide (PI-2610, HD Microsystems) was chosen as the electrode substrate and dielectric for strength (tensile strength of 350 MPa, vs 60 MPa for SU-8, 70 MPa for parylene-C), longevity, and biocompatibility [55].

See Figure 3 for an illustration of the fabrication process. This begins with new 6” p-type test grade silicon wafers. The native oxide on the silicon surface is kept intact in order to prevent excessive adhesion of the polyimide film to the substrate. A polyimide film of 2 *µm* is spin coated on the wafers (30s @ 500rpm, 0s @ 0rpm, 45s @ 3000rpm) and placed in a vacuum oven shortly after to cure in a nitrogen environment at a maximum temperature of 325 °C and pressure of 300 torr. Once the curing oven has cooled down, a bilayer of 1 *µm* LOR-5A resist and 1 *µm* i-line photoresist (OiR906-12) is lithographically patterned in preparation for the lift off process. Next, desalt is performed on polyimide by immersing the wafers in a 1:20 dilution of hydrochloric acid in DI water for 2.5 minutes followed by thorough rinsing in DI water and blow drying. Next, a platinum film 130 nm thick is deposited by electron beam evaporation. In order to obtain good adhesion between the metal and the polymer, a base pressure 9e-7 torr was achieved prior to the evaporation process. In addition, to prevent excessive heating of the photoresist two strategies are employed: first, ground-flat custom aluminum heat sinks are placed directly on the back surface of the wafers, with a thin layer of vacuum oil between them to provide thermal contact; second, the initial 30 nm of platinum is deposited at a high rate in order to reduce the amount of infrared energy absorbed by the photoresist. After metal deposition, lift off is conducted in a bath of PRS-3000 followed by generous rinsing in DI water. Our preferred metallization scheme was Pt, but in some occasions an alternative Cr/Au metallization scheme was successfully used.

**Figure 3:**
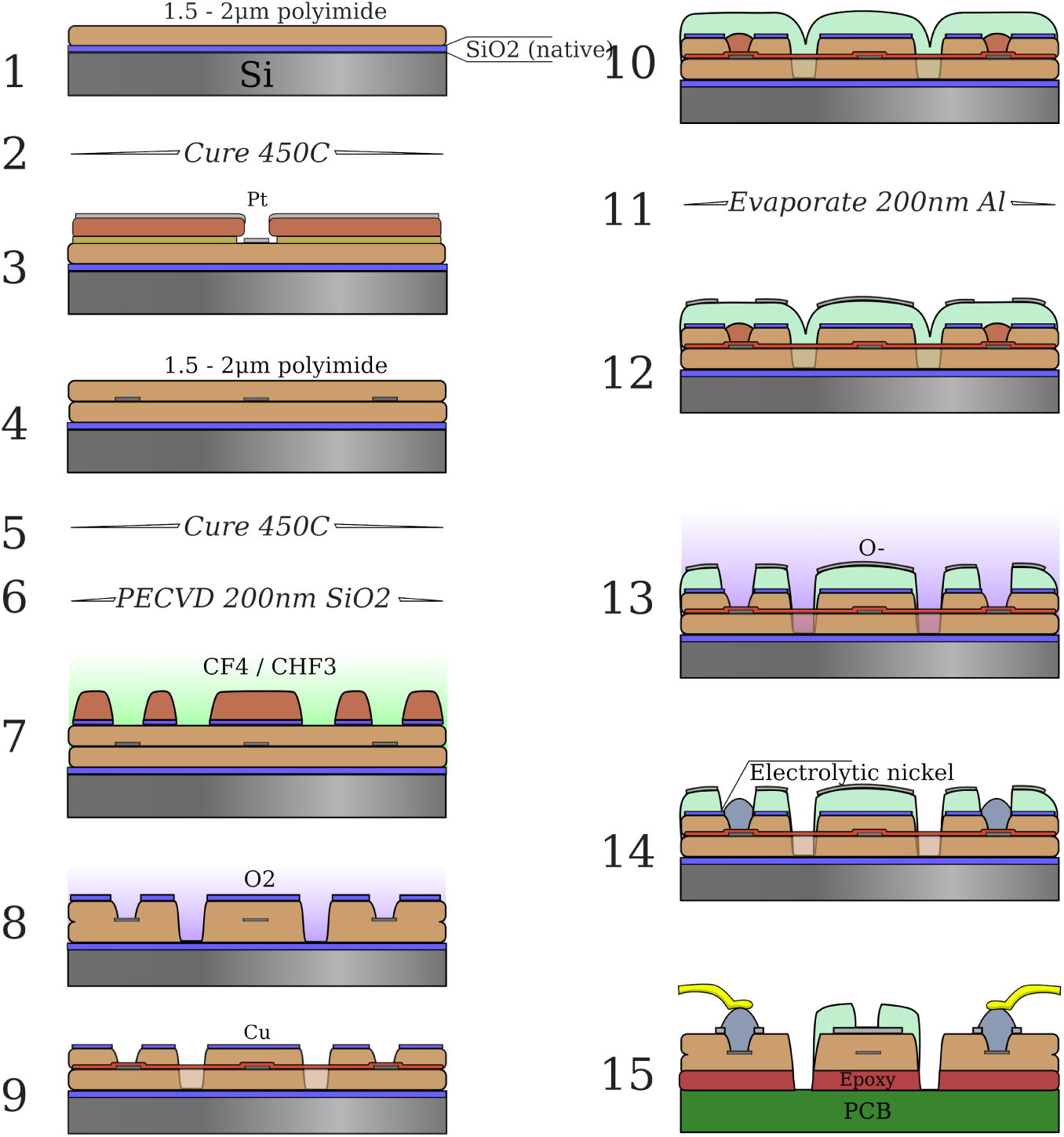
**1**. 1.5-2 *µm* polyimide is spin coated over a clean, native-oxide, 6” wafer, and **2** cured at 325 °C. **3**. Lift-off photoresist is patterned for the conductor traces, and 130 nm Pt is evaporated. 4. A second layer of polyimide is spin coated, and cured 5 at 450 °C. **6**. 200 nm SiO2 hardmask deposited via plasma ECR and **7** patterned via RIE. **8**. Device outline patterned using oxygen plasma. **9**. 400 nm Cu evaporated for the electroplating current busses and patterned via *FeCl*_3_. **10**. 5-6 *µm* parylene deposited via the Gorham process. **11**. 200 nm Al hardmask evaporated and **12** patterned. **13**. Parylene etched in oxygen plasma and Al is stripped. **14**. Ni is electroplated to form bondpads. 15. Device released and ultrasonically wirebonded using Al wire.

After a 10 minutes dehydration bake at 120 °C in an oven, a second layer of polyimide is spin coated with the same parameters as the first one and cured at a maximum temperature of 450 °C in a nitrogen environment at 300 torr (In the event of Cr/Au metallization, curing of second polyimide layer was done in two stages, first cure at 250 °C, cool down and then cure at 450 °C, in order to achieve optimal mechanical properties of the polymer films). A hard mask of silicon dioxide was used to pattern the polyimide. For this, a 200 nm thick *SiO*_2_ film was deposited by plasma enhanced chemical vapor deposition (Plasma Quest ECR PECVD System), followed by patterning via 2 *µm* g-line photoresist (OCG825-35CS). The oxide layer was etched by reactive ion etching (RIE, Oxide Rainbow Etcher, Lam research) and the polyimide layer was then etched by RIE in oxygen plasma (Plasma-Thermal Parallel Plate Plasma Etcher). This oxide layer is of critical importance: it serves as the weak, releasable adhesion layer between the parylene and polyimide, which allows electrodes to peel off without breaking the needle, and without prematurely falling off, e.g. when removing the electrodes from the carrier wafer.

The next step is to add the copper bus, which serves as a low resistance path for electroplating. 400 nm of copper are deposited on the entire wafer surface and patterned by *FeCl*_3_ wet etch using a g-line photoresist mask. Copper is removed from the entire electrode and bondpad area and only left on the perimeter of the wafer and along a wide trace in between devices as shown in figure 2C. The copper bus connects to each individual Pt bondpad through a 10 *µm* bridge in the Pt layer embedded in the polyimide. Once the copper etch is completed, a 5-6 *µm* parylene-C film is deposited (Parylene deposition system 2010). A hard mask of 200 nm aluminum is deposited on top of the parylene by electron beam evaporation and subsequently overlaid with lithographically patterned g-line photophoresist. The exposed aluminum is then removed by wet etch in aluminum etchant (Transese, inc.) and the now exposed parylene is etched in oxygen plasma. After parylene etch, the aluminum hard mask is fully removed by wet etch, as it hinders the passage of water vapor through the film stack, thereby making the probes harder to remove from the carrier wafer.

The last step is to electrochemically deposit nickel on the platinum bondpads. Standard photolithography with g-line photoresist is used to protect the regions of the wafer that do not require nickel plating. Using a custom jig, the wafer is immersed in Krohn Bright Nickel electroplating solution and connected to the negative terminal of a DC power supply. The positive terminal of the power supply is connected to a pure nickel anode also immersed in the solution. Electroplating is done for 15 to 20 minutes at 1.6 V. The photoresist is removed in an acetone bath followed by IPA and water rinsing. If Cr/Au metallization was used, a 5 second dip in chrome etchant is used to remove any chrome that may have diffused through the gold film during the high temperature cure.

Devices are released from the wafer by immersing the wafer in water and peeling off the polyimide devices carefully with sharp tweezers. During peel off, the Pt bridges used for electroplating break eliminating the electrical short between pads. Devices are transferred from the wafer to individual glass slides and then attached to custom PCBs. The latter is done by applying small drops of epoxy (Epotek 353ND) on the PCB and then laying down the thin film device on it and allowing capillary action to distribute the epoxy within the interface. A very thin and bubble free epoxy interface is important to facilitate wirebonding. The nickel bumps are then connected to the gold pads on the PCB by aluminum wedge bonding. Lastly, marine grade epoxy is used to encapsulate the bondpad area and the wirebonds.

### 2.3. Sewing machine and the insertion process

#### 2.3.1. Overview of the sewing machine device

Figure 4 shows the complete sewing machine robot. It is composed of several key parts: an inserter head, an imaging head, and a set of linear drives for positioning of both relative to the stereotax and animal. The imaging head (A) captures high-resolution images of the cortical surface through long-working distance objective (L), to be used for micro-durotomy via the laser (Q) and focusing objective (M). The imaging head is mounted rigidly to the primary XYZ axes, which serves as the reference coordinate frame. Following this, the primary axes (O,P) move the imaging head out of the way, the electrode cartridge (H) is lowered to be posterior the craniotomy, and the inserter head (C-F,J) proceeds to peel and insert electrodes. The inserter head is actuated via three secondary XYZ linear translation stages, allowing dependent and relative motion to the primary axes. Details for these steps are in the next two sections.

**Figure 4:**
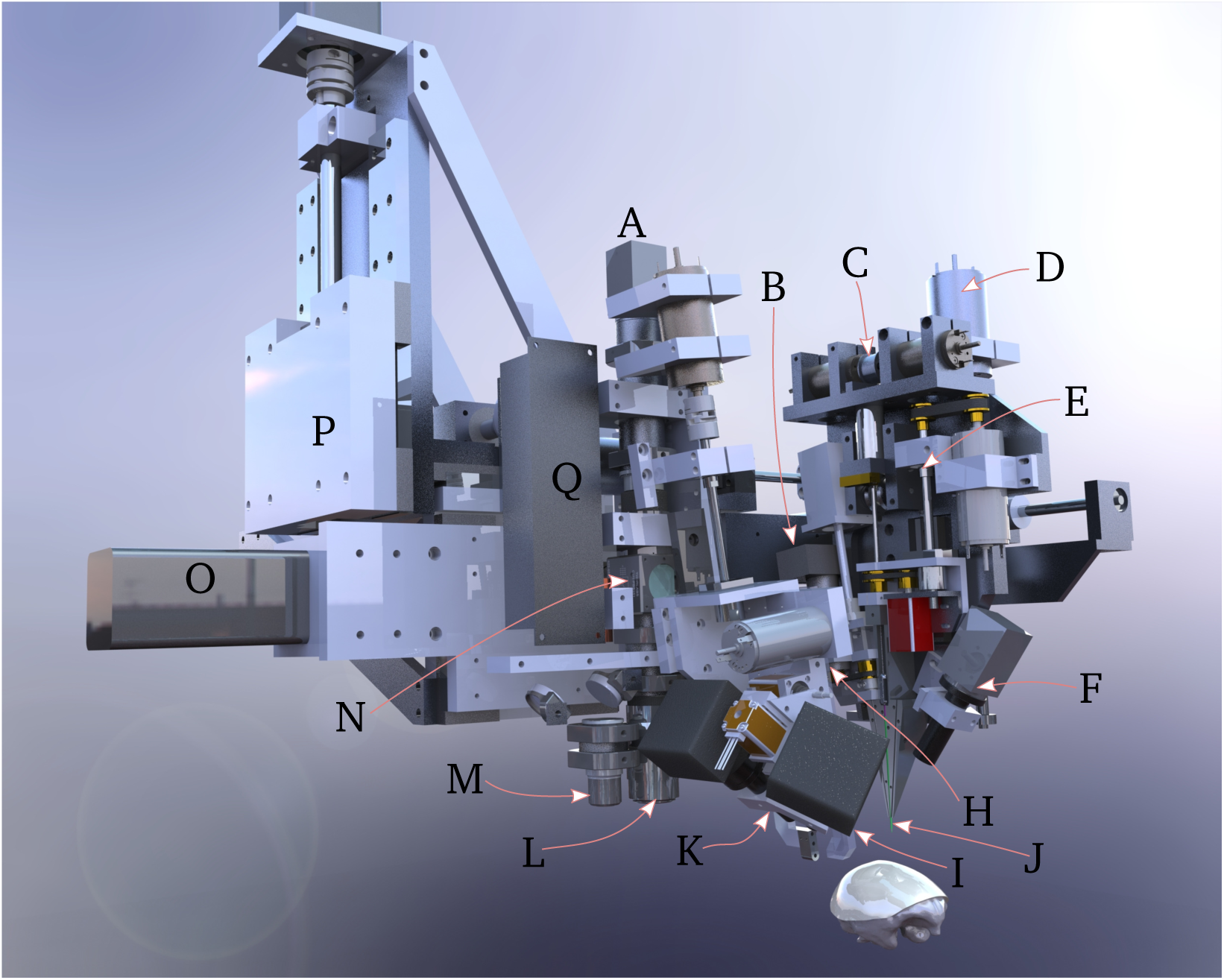
Inserter system overview. The scull of a rhesus macaque is shown for scale. **A**. Targeting camera stack. The system uses a 1/1.2” format 2.3MP monochrome camera and a standard f=200 mm tube lens. **B**. AC servomotor and linear stepper motor for pincher *θ* and *Z* control. **C**. Ballistic retraction mechanism. **D**. Motor for secondary Z axes. The secondary cartesian axes are driven by 150W coreless motors, and are mechanically chained from the primary cartesian axes. **E**. Ballscrew for high-precision actuation of needle depth during insertion. **F**. 1x oblique microscope and 1/1.8” 5MP monochrome camera for monitoring electrode peel-off and insertion. **H**. Retractable electrode cartridge stage. This whole assembly moves out of the way when imaging the brain surface and targeting the insertion sites. In the foreground is the motor for rapid backlight retraction (silver cylinder), and the 90° and 45° needle-electrode targeting cameras (black squares). **I**. Retractable backlight. During needle-electrode targeting, the backlight is behind the needle to regularize illumination, independent of surgical lighting. A 405 nm LED and diffuser is mounted to this backlight arm; polyimide is absorptive at this wavelength, yielding high-contrast images of the electrode loops. **J**. Needle cartridge, green, composed of four telescoping cannula. Pincher is located at an angle to the left of the needle cartridge. Additional lines here include air puff and saline squirt. **K**. Electrode targeting mount. This has two hybrid lead screw-stepper motors for controlling focus and scan of the two cameras across the electrode cartridge knife-edge. **L**. 2x long-working distance, large FOV objective for imaging the brain surface. In combination with the targeting camera stack *A*, this yields an image field of 5.4 × 4 mm. M. High power 5x objective for laser micro-durotomy. **N**. Polarizing beamsplitter for inline illumination and glare rejection. **O**. Primary X axis. These axes are driven by 400W AC servomotors with 4k encoders coupled to C5 grade ballscrews. **P**. Primary Z axis mounting point. Primary Y axis is behind laser. **Q**. Microdurotomy laser.

#### 2.3.2 Imaging and micro-durotomy

The first stage of the automated insertion process uses the imaging head to accomplish two tasks: identification of target insertion sites and laser-ablation to create micro-durotomies at each site. The targeting phase is straightforward: the surgeon manually selects implantation sites via the targeting camera, which are then translated via a calibrated offset to robot coordinates. Image coordinates determine X and Y, while Z is controlled by focusing over the curved brain surface. ‡

The next step is to perform microdurtomies at the insertion site. There are two motivations for this step in place of macro-durotomies. First, manual durotomies can result in bruising of the cortical surface. Second, in earlier studies, we have observed post-mortem that increased intracranial pressure sometimes leads to significant cerebral hernation through the macro-durotomy. Ultraviolet light, as is used in LASIK, is preferred for controlled ablation of tissue; however given the difficulty of procuring or producing a UV laser (e.g. either the 3rd or 4th harmonic of Nd:YAG), the potential for absorption in protein-loaded CSF, and then challenge of integrating such a laser with the robot, we chose to take a different approach: staining the dura with a red dye and then using a green laser to perform tissue ablation.

**Figure 5:**
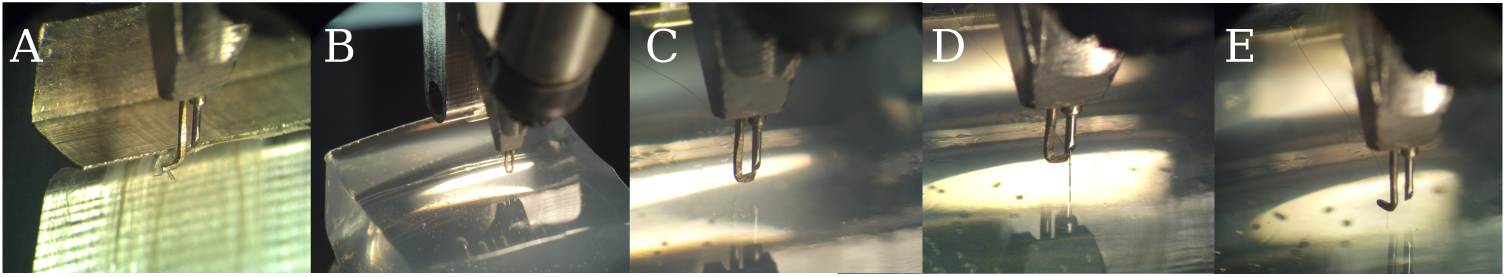
**A**. Needle is threaded through the electrode loop, which is exposed past the knife-edge of the electrode cartridge. **B**. The needle-pincher peels the electrode off the parylene backing sheet which is glued (Zap thin wicking cyanoacrylate) to the electrode cartridge. In this figure, electrodes are being inserted into a 0.6% w/v agarose tissue proxy. The pincher is used to prevent the electrode from falling off the needle, and to support the needle and electrode so the needle does not bend during rapid peel-off. **C**. The needle and electrode are moved over the target site, and the pincher is slightly retracted to act as a pulley to keep the electrode from slicing laterally into the brain. **D**. The needle is lowered to drive the electrode to its anatomical target. **E**. The needle is retracted, the pincher moves out of the way, and the process repeats.

The dura is first stained by applying 400-500 *µl* of saturated solution of erythrosin-B in 0.9 *N* saline on the surface. Erythrosin B is a food-grade polar dye, which sticks readily to protein; it also offers peak absorption at ∼ 524 nm, which is close to the emission spectra of available compact frequency-doubled Q-switched lasers. Staining is applied for 4 minutes or more, as the more highly stained the dura is, the greater the proportion of laser energy absorbed there, and the less energy penetrates the brain. Here we have used a 527 nm KTP-doubled Q-switched Nd:YLF laser (pulse energy 230 *µJ*, 1 kHz, pulse width 15 ns) for micro-durotomy; as few as 3-4 pulses are sufficient to penetrate the dura. The green laser had a tendency to burst small blood vessels, as they also strongly absorb green light. To minimize this effect, a 405nm, 1W diode laser was coupled into the beampath via a dichroic; this wavelength is near the peak hemoglobin absorption, and served to cauterize any resultant small bleeds.

**Figure 6:**
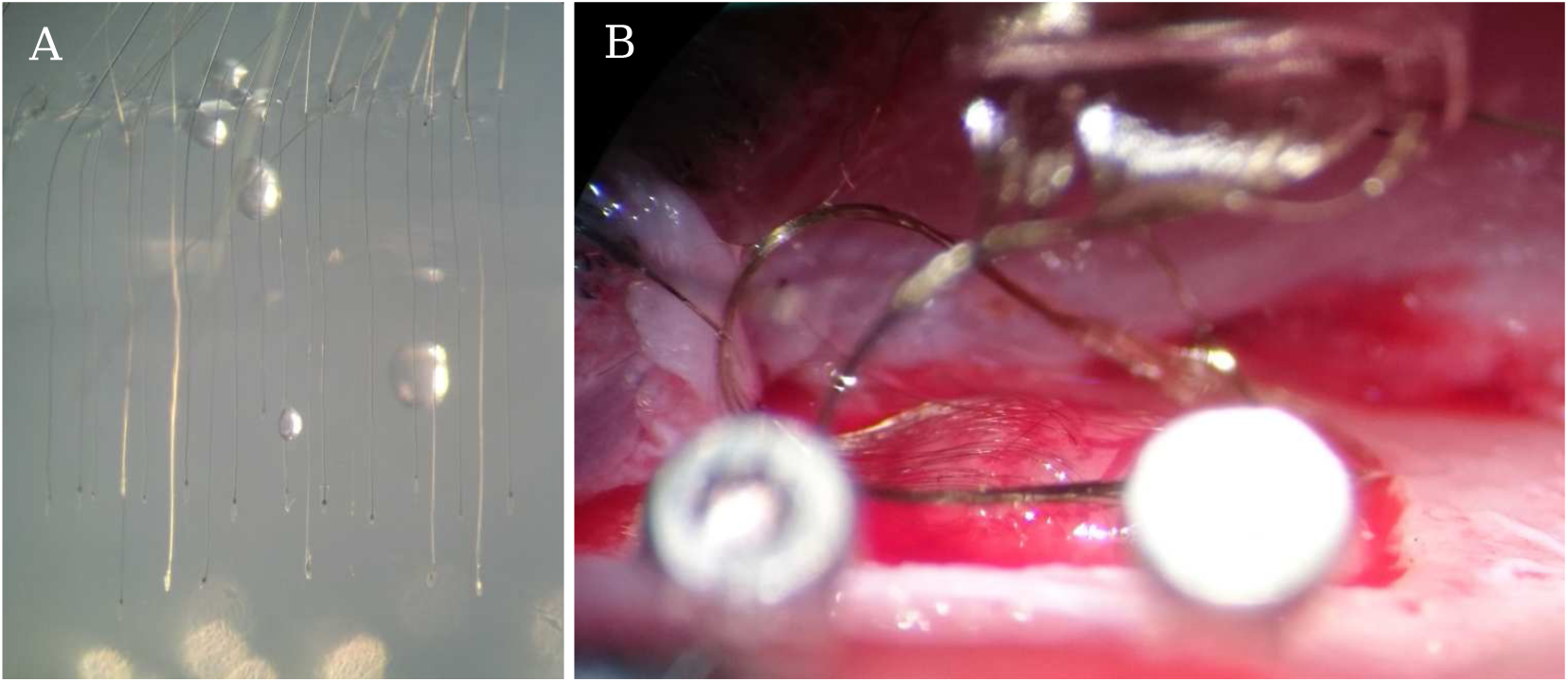
Examples of insertions. **A**. Early result showing 20 insertions into 0.6% w/v agarose tissue proxy. Electrodes were in two rows; bubbles are from the halogen lights used in the test. **B**. Image showing 42 electrodes inserted into the somatosensory cortex of a rat (#10). One needle was used.

The surface imaging camera uses 590 nm illumination from a set of LEDs (not shown in figure 4), which is within the hemoglobin absorption band, but outside the erythrosin-B absorption – hence the vessels remain clearly visible even in the presence of the dye. Polarization is used to reduce glare from saline meniscus; illumination polarization is perpendicular to light transmitted by the polarizing beamsplitter in the targeting camera stack, hence most imaged photons are scattered. See supplemental figure 11 for a depiction of the imaging system.

#### 2.3.3. Automated insertion

Once insertion targeting and microdurotomies are complete, the imaging head moves out of the way, the electrode cartridge stage is lowered, and electrodes are inserted. The insertion process is illustrated in Figure 5.

First, needle step depth and orientation is calibrated so that the step is slightly out of the 34 gauge cannula, and oriented facing away from the knife-edge, so that it can optimally engage with the electrode loops; see supplemental Figure 12 for an image of the needle step. This calibration needs to be done once per needle; the needle cartridge itself may be independently sterilized and replaced before and during a surgery. Then, for each electrode, the needle is threaded through the loop. The control software (described in the supplemental section 8.1 *Software*) positions the needle within the view of the two electrode targeting cameras (Figure 4 I,K) and uses computer vision to guide it through the loop. Next, the pincher (a 90° bent piece of 50 *µm W* wire) is rotated around to pinch the electrode to the lower needle cannula (34 gauge, 150*µm* outer diameter). This is an important step, as it buffers the needle from the stress of peel-off – the set of needle-electrode motions that separates the polyimide electrodes from the parylene backing sheet – thereby keeping the needle from bending. As such, the degree adhesion between the parylene and polimide is critical during peel-off, as described in section

### 2.2 *Electrodes* above

The peel-off maneuver leaves ∼ 20*mm* of electrode lead freely floating in air so that insertion can proceed under minimal tension. The inserter head then moves to the target site, whose coordinates are determined in the imaging phase above. The pincher moves slightly out of the way to act as a pulley for the electrodes as they are inserted, ensuring the threads followed the needle path, rather than cutting laterally into the tissue. The needle then drives the electrode into the brain through the micro-durotomy. Then bursts of air or saline from embedded cannulae (4J) may be used to lay the electrode threads onto the surface of the dura. The needle then retracts with high acceleration and jerk using the ballistic retraction mechanism (Supplemental Section 8.3). This serves to keep the electrodes from coming out with the needle, or from buckling and being driven laterally into tissue as the needle retracts, both which are otherwise significant problems. With the needle retracted, the inserter head moves back over the electrode cartridge to target the subsequent loop in the array.

### 2.4. In vivo demonstration

For an in-vivo demonstration of this system, we used adult male long-evans rats, 300-500g. All procedures were performed with approval of the University of California, San Francisco Institutional Animal Care and Use Committee and were compliant with the Guide for the Care and Use of Laboratory Animals.

#### 2.4.1. Surgical methods

First, rats are anesthetized with 4% isoflurane in oxygen, their skin cleaned in betadine and sterile saline, and an incision is made along the midline to expose the cranium. After incision, isoflurane is lowered to ∼2%. Temperature is maintained at 38 °C with a heating pad. The cranium is scraped and cleaned with 3% hydrogen peroxide, and a *<*5 mm (A-P) × 4mm (M-L) craniotomy is marked on the dorsal surface, corresponding to somatosensory-motor cortices. Six to eight bone screw holes are piloted via a FG1 dental drill, and bone screws (M1.2 thread-rolling SS316 screws, 4mm length, McMaster-Carr) installed and secured with a thin layer of MetaBond luting cement. The craniotomy is then completed and the dura was flushed with saline prior imaging, micro-durotomy, and insertion.

Following insertion, the probe shanks are gathered into a small bundle with Dow-Corning silicone gel, 3-4680, which also serves to protect the brain. Then the connector PCB is positioned along the sagittal plane, and everything is sealed and affixed with PMMA dental acrylic (Lang Jet Denture repair). Animals are then given a bolus of saline, taken off isoflurane, and once fully recovered, given buprenex (0.03 mg/kg) and meloxicam (1 mg/kg) SQ. Post-op analgesic and antibiotic treatment was continued for 3 days.

#### 2.4.2. Electrophysiological recordings

All recordings were done with our Tucker-Davis Technologies RZ2 attached to a PZ2 amplifier/digitizer and 64-channel ZIF-clip active headstages. To mate to the ZIF-clip, we designed a custom passive wirebond adapter PCB, as mentioned in Section 2.2. Impedance measurement was performed using a 32-channel TDT IZ2 stimulator using manufacturer supplied software.

#### 2.4.3. Histology

Animals were deeply anesthetized with an overdose of Euthasol and perfused intracardially with heparinized phosphate-buffered saline (PBS), followed by fixation in 4% paraformaldehyde in PBS, pH 7.4. In animals in which the electrode cap came off (or was removed) prior perfusion (#13 - 19d, #14 - 14d, #11 - 39d)) brains were extracted on the same day and postfixed in fixative overnight. For animals with the head mount intact (#12 - 29d, #16 - 72d, #17 - 25d) brain extraction was performed in several steps so that that implanted electrodes were not pulled out or displaced during the procedure. First, brains were partially exposed on a ventral side and placed in fixative overnight at 4 °C. In a following day the remaining skull bone, acrylic and other parts of implant were carefully removed under a dissecting microscope using a dental drill and microsurgical tools; electrode threads were cut about 1 mm above the brain surface with microscissors leaving inserted parts in the brain undisturbed. Extracted brains with threads were immersed in 30% sucrose for 5-6 days at 4 °C and frozen at −20 °C for cryosectioning. Coronal 50-80 *µm* thick serial sections were cut on a cryostat (Microm, Thermo Fisher Scientific), and collected in 12-well plates filled with PBS.

To reveal changes in neuronal tissue in response to the implantation procedure, sections representing surgical areas were stained using a routine Nissl staining (0.1% Cresyl Violet). To show changes in glial cells (astrocytes) sections were immunolabeled for GFAP (glial fibrillary acidic protein). Briefly, free-floating sections containing regions of interest with flex electrodes were permeabilized with 50% ethanol for 10 min and rinsed in PBS. Sections were then blocked with 10% normal donkey serum in PBS for 30 min and incubated for 48 hrs at 4 °C on an orbital shaker with a primary anti-GFAP antibody, (mouse mAb Sigma-Aldrich, 1:1,000) diluted in PBS containing 0.05% Triton X-100. Next, sections were washed in PBS, incubated with 2% normal donkey serum for 10 min, and incubated for 4 hrs with the secondary Alexa Fluor 488-labeled donkey anti-mouse antibody (1: 300) for immunofluorescence. For immunoperoxidase staining, sections were initially incubated in 3% hydrogen peroxide/PBS for 10 min to quench endogenous peroxidase activity, then processed as above, until the secondary antibody step, which was the incubation in biotinylated donkey anti-mouse antibody (1:500, Jackson Immunoresearch) for 4 hours, followed by rinses in PBS and incubation in ExtrAvidin (Sigma-Aldrich, 1:5,000) for 4 hours. Peroxidase was detected using diaminobenzidine (Sigma-Aldrich). Sections were rinsed, mounted on gelatin-coated slides, air-dried, dehydrated in graded alcohols, cleared in xylene and coverslipped with D.P.X. mounting media (Sigma-Aldrich). After immunofluorescence, sections were rinsed in PBS and coverslipped using ProLong Gold Antifade mountant medium (Thermo Fisher Scientific). Images were acquired using laser confocal microscope LSM510 Meta (Carl Zeiss Microscopy, LLC) and Nikon Eclipse 90i wide-field microscope (Nikon Instruments Inc)

## 3. Results

We have repeatedly and reliably implanted our thin, flexible electrodes in both agarose tissue proxy and the brains of rats (Figure 6). In Figure 6B, note that the individual threads form a single “rope” that loops up to the PCB and connector assembly. This formation results from capillary action. Provided the free length of electrode is not too long and is constrained during peel-off and insertion, capillary action allows the electrodes to remain both reasonably well organized and mechanically isolated when affixing the connector to the scull.

Regarding reliability, insertion into agar is consistent and repeatable; reliability in-vivo is limited by surgical factors, mainly: blood occluding the surgical site, blood clogging the needle cannula, microdurotomy quality, needle durability, and implant durability. More explicitly: avoidance of blood vessels keeps the sugical site clean, which allows for accurate targeting of subsequent insertions. If a vessel is inadvertently hit, blood tends to spread to form a subdermal hematoma, occluding future insertions; some of this blood will leak to the surface, which can wick up the needle guide-cannula and clot, potentially causing the needle to seize. Likewise, if a microdurotomy is too deep, subsurface capillaries may be ruptured, again occluding the surface and possibly contaminating the needle cannula; if it is too shallow, sufficient dura may remain to buckle the needle, forcing replacement. Nonetheless, most surgeries were done with care and with one needle. Finally, as with many other approaches, implant durability is limited by the bone-screw’s attachment on the growing and flexible rat cranium; this failure mode is shared with other approaches.

Because the whole insertion process is automated and the robot can move quickly, per-thread insertion times were quite short. Indeed, other than the needle insertion, which is performed at 0.1-2 mm/s, robot movements are rapid, up to 100 mm/s. This allows for the per-thread insertion cycle times of less than 9 seconds (supplementary video 1). Such rapid insertion allows for large numbers of electrodes to be inserted in a practical surgical timeframe.

### 3. Histology

In animals in which electrodes were removed prior fixation, examination of sections stained with Cresyl Violet and GFAP immunohistochemistry has revealed multiple penetration lesions (Figure 7 A,B). In vivo insertions resulted in a different pattern from what might be the expected after the agarose insertion of Figure 6. Although most of the penetration lesions were generally in downward direction, these varied significantly in their size, shape and exact orientation (arrowheads in Figure 7, tracks #1-7 in 8 A). Some spanned all cortical layers and could be followed into the white matter and subcortical structures, whereas others were more oblique and ended more superficially in depth (not shown). The majority of lesions were wider near the cortical surface and narrower at the deeper end becoming to be about 50 *µm* wide. Some lesions were larger, up to 1-2 mm wide (Figure 7A,B, lesion #7). Furthermore, several adjacent lesions were seen as merged at the top forming one large area of the eroded layers I-II (asterisks in Figure 7 A,C). The tissue in subcortical penetrations such as in the dorsal hippocampus were less affected (not shown). To estimate more accurate position of inserted threads and especially the location of the recording sites, brains in two animals were extracted and cryosectioned with electrodes remaining in situ (Figure 7 C,D). Microscopic examination has confirmed that tissue sections had the ability to retain fragments of implanted electrodes without displacement even after cryosectioning, thawing and immunostaining. Most electrodes had recording sites in deeper layers V and VI which as had less tissue damage and consistently showed more normal morphology than upper layers (Figure 7 C); as shown in Figure 7, D1-D4 recording parts of the flex electrodes in layer V were surrounded by neuronal perikarya with normal appearance. Few electrodes were found to be inserted very obliquely with recording sites in layers II and III (e.g. Figure 8 A). All lesions had little or no surviving neuronal cells at its core and increased density of non-neuronal cell (e.g. densely filled core of the lesion #7 as shown in Figure 7 A); these were morphologically identified as granulocytes (neutrophils), monocytes (macrophages), microglial and astrocytic cells (not shown). GFAP (an astrocytic marker) was upregulated in all examined animals, and expressed at higher levels at the insertion sites (Figures 7 B, 8 A), and in upper layers (8 A). High resolution microscopy revealed dense plexuses of GFAP-positive astrocytes that were lining up the space around the implant cavity (Fig 8 A); astrocytic processes were also in a direct contact with the implant material (Fig 8 D).

**Figure 7:**
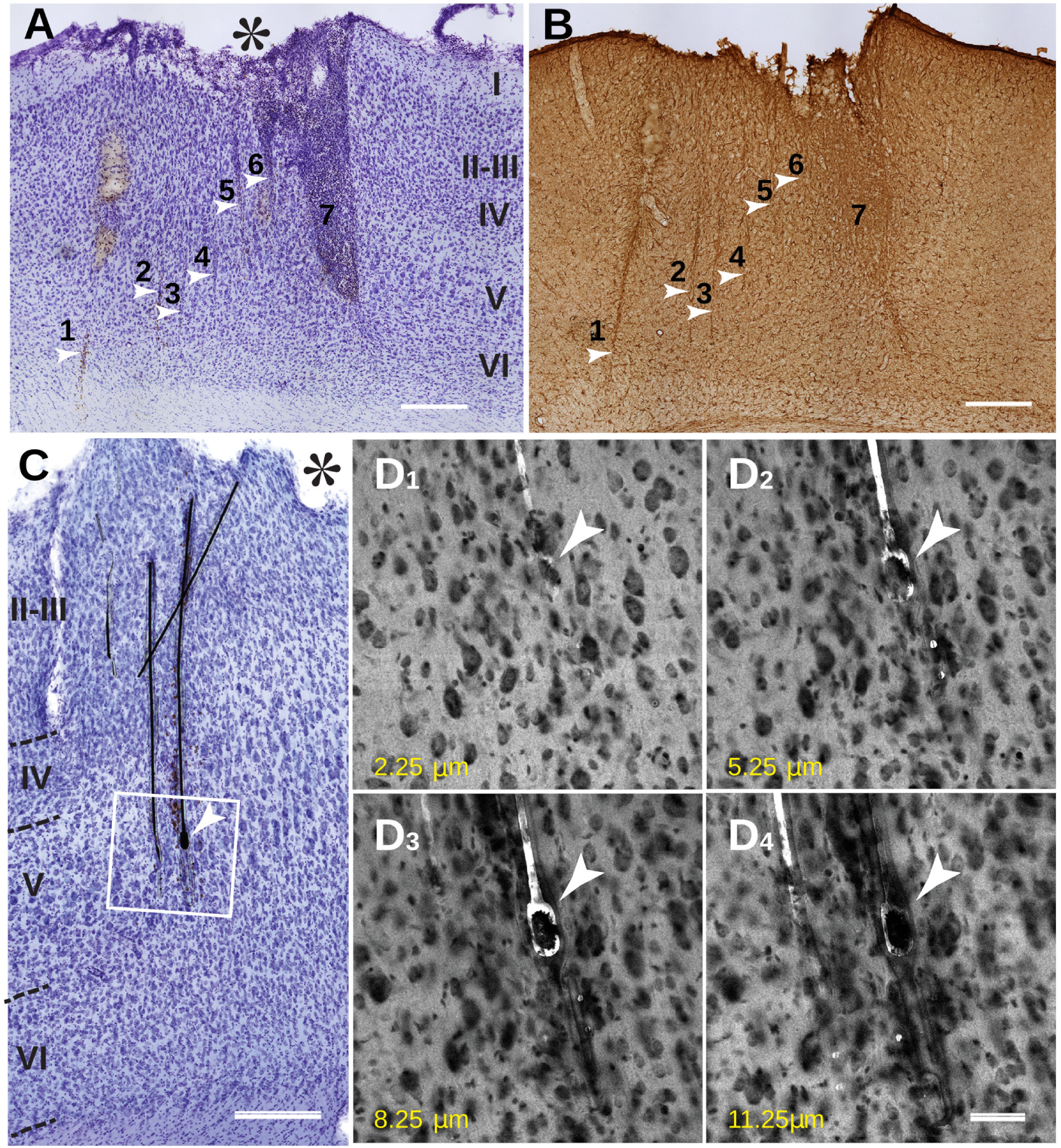
Histological changes in the cortex after sewing. A,B show cortical tissue response in an animal implanted for 2 weeks. C,D show the response after 10 weeks of implantation. **A**. Cresyl Violet (Nissl) staining show presence of insertion lesions (arrowheads), numbered 1-7; roman numerals I-VI indicate cortical layers; Note a much large lesion # 7 filled with non-neuronal cells, also lesions 5, 6, and 7 and likely more have merged at the top and formed a large necrotic cavity (asterisk). **B**. Same as in A lesions are also seen with GFAP Immunoperoxidase (#1-7, arrowheads) on the adjacent section. Scale bar: A, B 500 *µm* **C**. Two vertically oriented and near parallel electrodes and one smaller fragment going across. The tissue damage was noticeably larger in superficial layers I-III (asterisk) and much lesser in layer V, which contained the recording site (arrowhead). Boxed area is scanned with a confocal microscope and shown in D1-4 to demonstrate that those electrodes were not merely stuck to the surface of the section but were inside the tissue. **D**.1-4: Consecutive confocal images from the boxed area representing a 3 *µm* step depth using confocal imaging in 488 nm laser and 80/20 neutral beamsplitter. This also reveals a tissue cavity that contains the implant as well as a more accurate relative position of neuronal cells bodies (seen as dark profiles) near the recording site (arrowhead on D1-D4). There are many neurons in within 10-20 *µm* of the immediate vicinity of the recording site Scale bar C, 250 *µm*, D1-D4, 50 *µm*.

**Figure 8:**
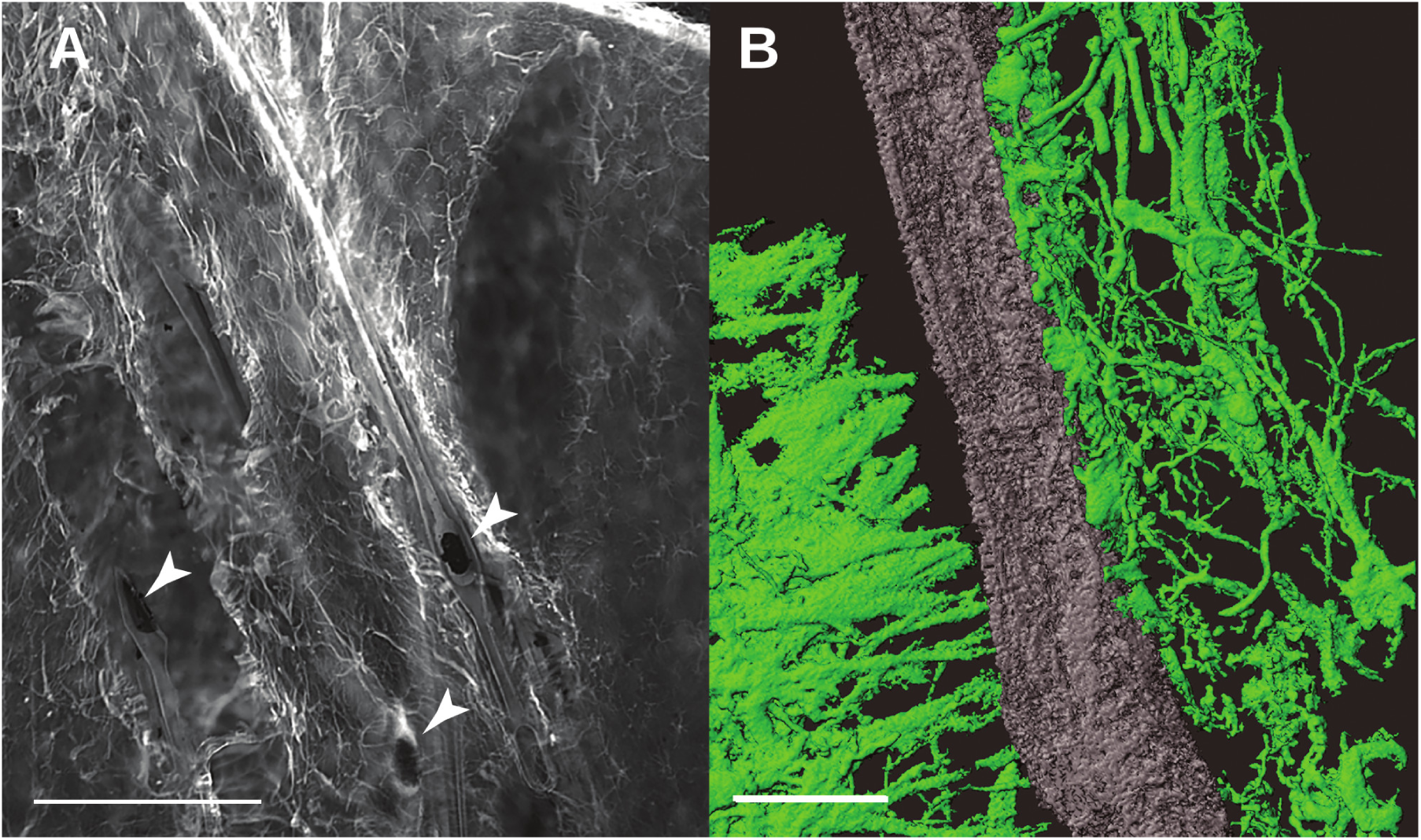
**A**. Area with implants (arrowheads) in the layer III showing upregulation of GFAP immunofluorescence, especially near the electrodes. Arrowheads point to electrode fragments with recording sites. **B**. 3D reconstruction of high resolution confocal images of the GFAP-immunoreactive processes from the same tissue as in A. Note areas of possible direct contact between the implant and astrocytic processes (arrows), and the relative preference of the astrocytes for one side of the electrodes. Unknown if this was the *SiO*_2_ (hardmask) coated side, or the polyimide side. Scale bars: A, 250 *µm*; B, 25 *µm*.

### 3.2. Electrophysiology

During the course of device testing and refinement, we performed extracellular recordings from four rats (rat #5, #13, #15, #16); other animals were used for device and insertion testing and refinement Data from rats #5 and #16 are shown in Figure 9. Rat #16 had 24 implanted electrodes that were recorded over 2 months. We observed single-unit action potentials on 39% of the implanted electrodes overall: 3 of 22 electrodes for rat #5; 2 of 13 electrodes for rat #13; 7 of 12 electrodes for rat #15; and 16 of 24 electrodes for rat #16. Rats #5, #13, and #15 had their implant fall off prematurely, and did not last as long as #16.

**Figure 9:**
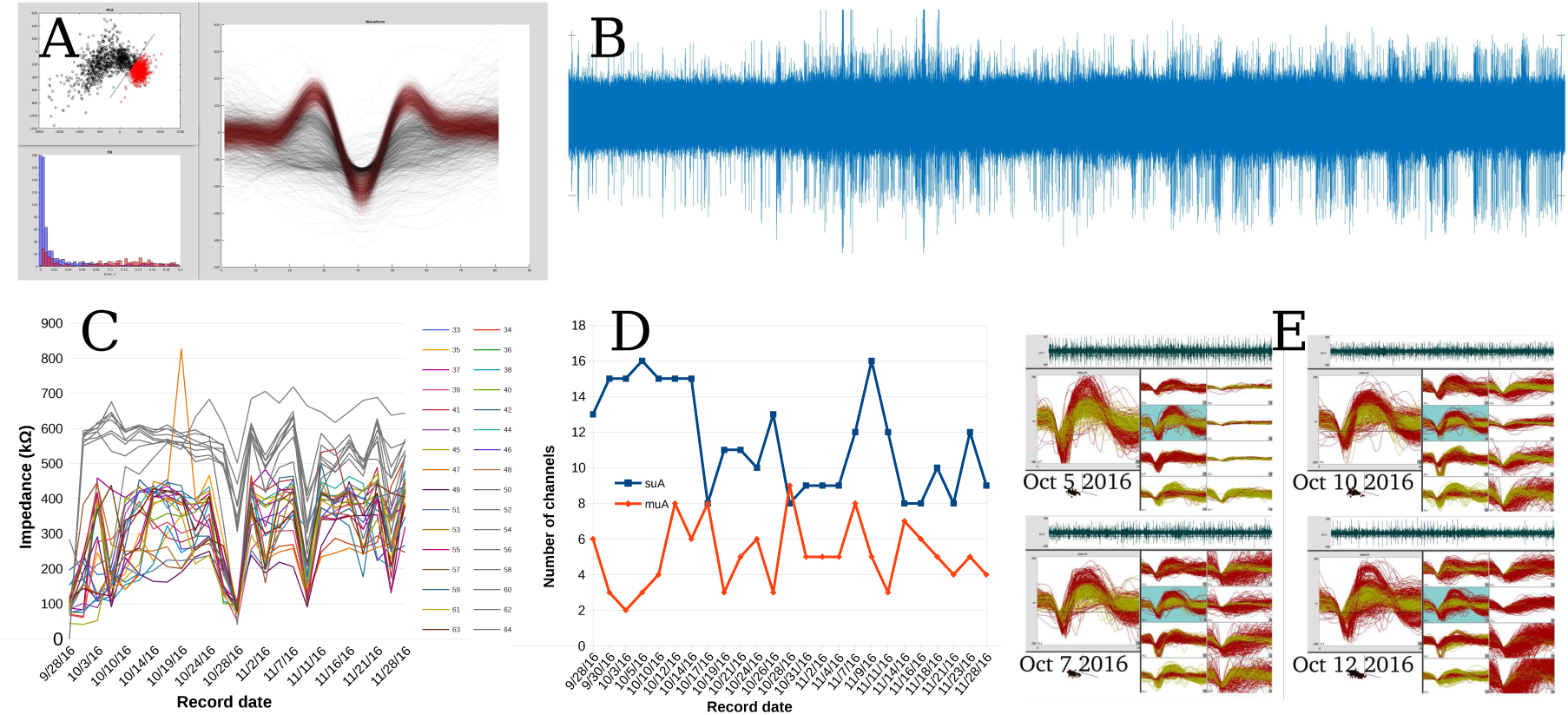
Representative electrophysiology. **A**. Example single-unit, showing PCA, voltage waveform, and inter-spike-interval distribution. This was the first ‘spike’ recorded with the system. **B**. Raw voltage trace from the same rat (#5). **C**. Evolution of electrode impedances in rat #16. Gray are unimplanted control electrodes, cut off above the brain. **D**. Recording quality over time. SUA = single unit activity, easily sortable. MUA = multi-unit activity, not sortable. **E**. Example waveforms over one week showing stability.

Figure 9 shows electrode impedance, as measured with our TDT system, from one rat (#16) implanted with 24 electrodes. Of the bank of 32, 8 electrodes (gray) were not implanted and cut off above the brain; these serve as a control. Note the TDT system cannot report or measure impedances *>* 1*M* Ω, so control impedances are qualitative. Implanted electrodes were plated down to ∼ 100*k*Ω prior implantation with platinum black (chloroplaticinc acid + lead acetate, −2*µA*, 60 s). Electrode impedance of the implanted electrodes gradually increased with time, consistent with the platinum black flaking off; impedance of the control electrodes gradually decreased, consistent with the connector, encapsulant epoxy, or polyimide absorbing water. Despite changes in impedance, recording quality was relatively constant over the implant period, with some waveforms stable up to a week.

## 4. Discussion

Cerebral cortex is a complex laminar structure consisting of dense layers of neuronal and non-neuronal cells and a thick layer of myelinated fibers underneath; it is covered with layers of protective membranous connective tissue; and finally, it contains a dense network of capillaries under considerable hydraulic pressure [56]. All these factors are contribute to cortical firmness and elasticity, hence it is no surprise that cortical insertion resulted in a heterogeneous pattern of implantation. The histological assessment has revealed that proximity to the surface was the most influential factor in the tissue response; the damage was strongest at the cortical layer I and II and dissipated in the deeper tissue, thus placing the priority for the sewing method more on reducing the amount of injury to upper cortical layers. It is not clear what exactly have caused such damage; on one hand, it might be caused by surgical procedures such as durotomy – or dimpling-related damage to pia and to capillary branches below the pia, which are much more numerous in layers I-III [34]. Another significant consequence of sewing on the tissue condition was presence of large penetrating lesions that were clearly associated with insertion process. It is also likely that the inserter needle can inflict a more significant injury by accidentally stabbing a larger blood vessel such as a penetrating arteriole on its way [57]. Overcoming these technical problems and improvements to the surgical techniques are underway. Because the damage to deeper cortical layers and subcortical structures appears minimal, it has also demonstrated a potential for using flexible electrodes in a chronic long-term recording experiments especially for subcortical structures.

Overall, sewing had shown three general sources of tissue damage:

1. The superficial damage possibly due to severing pial blood supply or other factors. This might have triggered neurodegeneration and cell death in upper cortical layers; it may also cause tissue swelling and herniation, thus unpredictably changing electrode placement.
2. Localized and contained penetrating lesions, occasionally large, up to 500 *µm* wide inflicted by the direct mechanical movement of the needle. This causes sparse cell loss and scarring in the vicinity of the track, and possibly reduced neuronal activity at the recording site.
3. Highly localized and contained reaction of the tissue to the presence of implant, such as tissue cavitation around the implant and local glial activation. There is no data to support that this may case neuronal loss, but it might affect electrical properties in the recording site.

It is very likely that the vasculature damage overwhelms all other factors, making lesions much larger by cutting blood supply and by triggering the migration of inflammatory cells such as neutrophils and monocytes (macrophages) into the area.

Nevertheless, as evident by a better preserved deeper layers and subcortical structures, small and flexible implants can benefit a long-term implantation outcome; however, the insertion procedure must be better guided at the microscale level to avoid impacting descending vessels. Future work can focus on improving this targeting, perhaps with OCT imaging and more degrees of freedom on the inserter head, as well as further miniaturizing the needle and electrodes, and lithographically patterning more electrode contacts per shank.

### 4.1. Conclusion

We have demonstrated that the sewing machine approach is effective in recording extracellular neural activity in rodents, and is worthy of further study and development.

## Supporting information

Supplemental Video: One insertion cycle of the rev 4 sewing machine.

## Acknowledgements

TLH would like to acknowledge the generous help of Eric R. Hanson in the machining and design of five different revisions of the inserter robot, Keenan M. Hanson for advice and help with metallurgy, Jasmine Amerasekera for fabricating the needles, David Piech for helpful discussions of surgical planning, Jason Chung, Hannah Joo, Daniel Liu and others from the Loren Frank lab for vital instruction and help with surgical technique, Lindsey Presson with surgical assistance, and Joseph Makin and Joseph O’Doherty for helpful scientific discussion.

This work was supported by DARPA Contract W911NF-15-2-0054 from the Biological Technologies Office (BTO). The views, opinions, and/or findings contained in this material are those of the authors and should not be interpreted as representing the official views or policies of the Department of Defense or the U.S. Government.

## 6. Author contributions

TLH, PNS and MM ideated the overall system. TLH, CAD, and MM designed the electrode fabrication process. TLH and CAD designed the lithography masks. CAD fabricated, released and bonded the electrodes. TLH designed and fabricated the inserter robots and wrote the control software, designed and fabricated the needle brazing system, performed surgery, and did electrophysiology. VK did all histology. All authors contributed to the manuscript.

**Figure 10:**
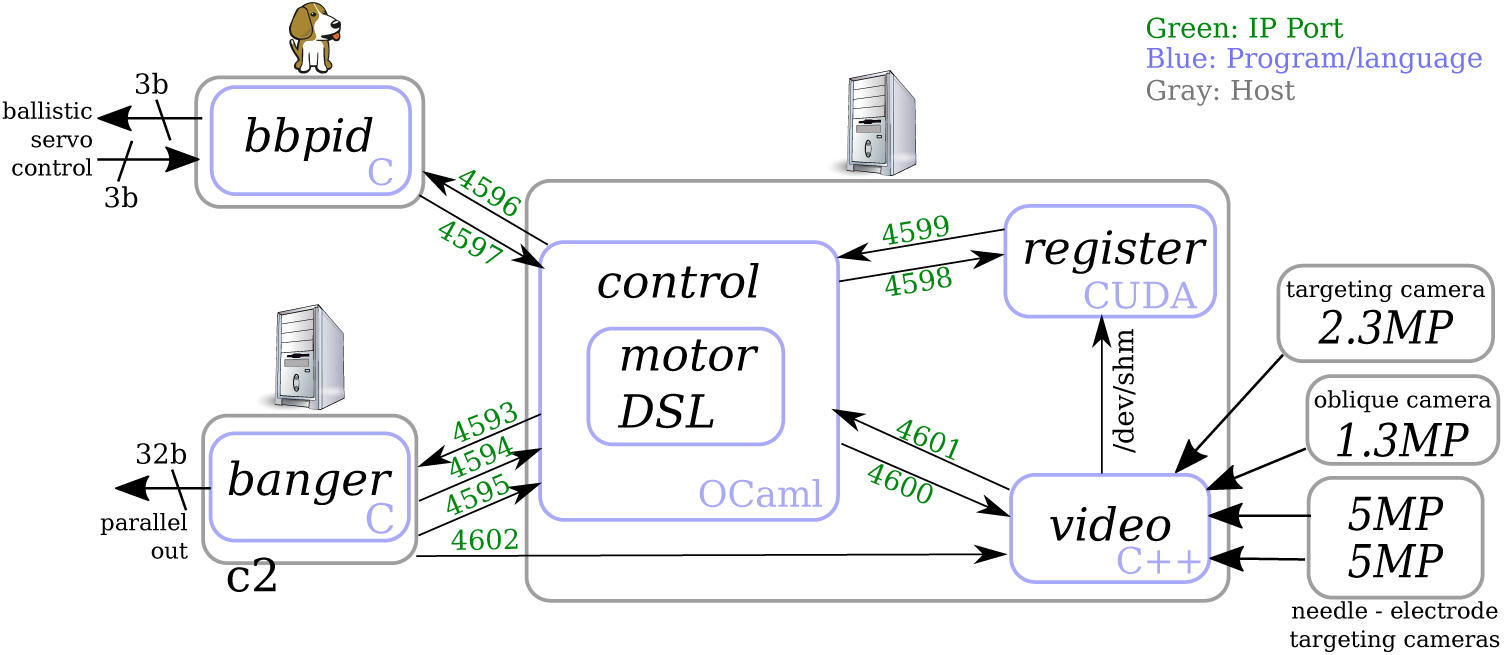
Software control system for the inserter robot. ‘bbpid’ refers to ‘Beagle bone proportional–integral–derivative’, which is a closed-loop controller for ballistic retraction.

**Figure 11:**
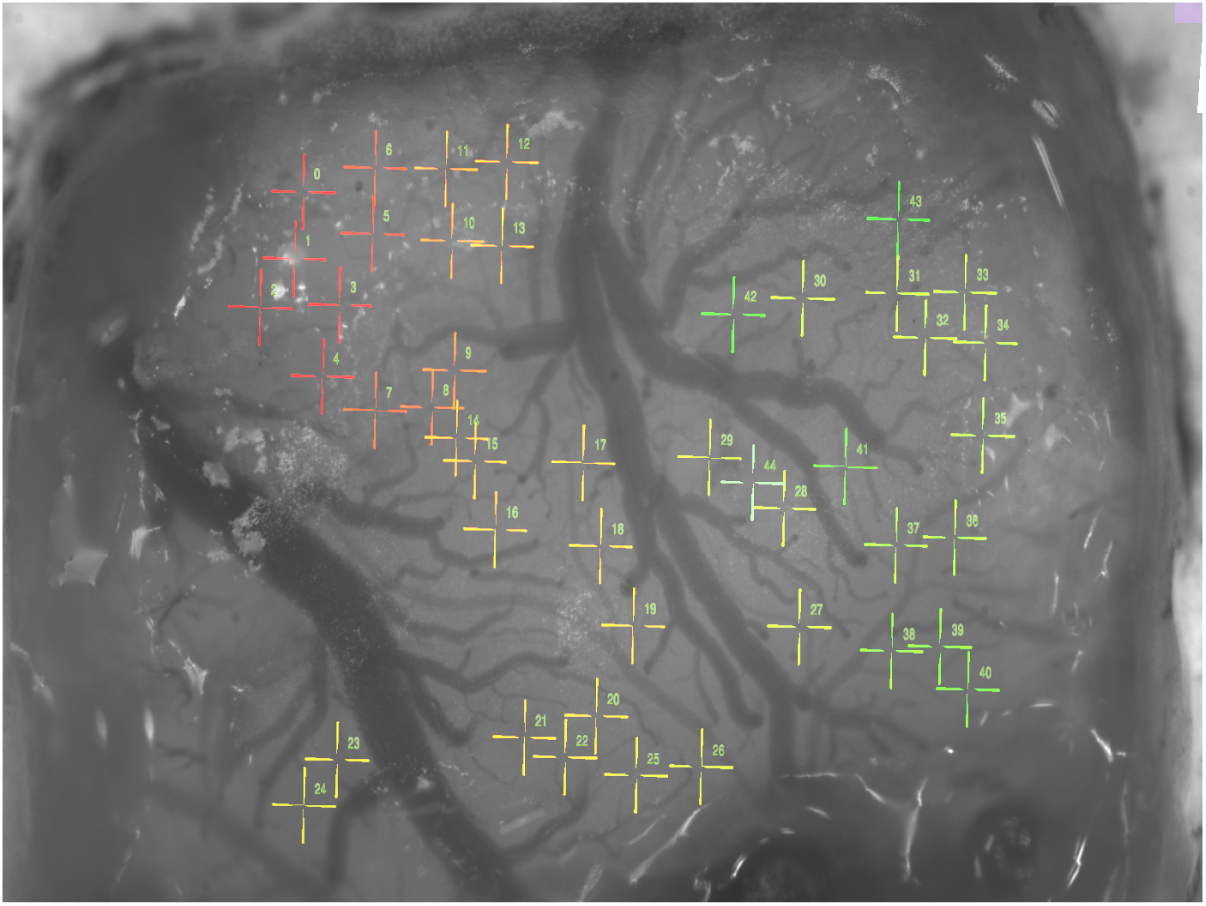
Image of the cortical surface, as captured using the targeting stack in Figure 4 A. Here the dura was stained with a saturated solution of Erythrosin-B in saline, and illuminatd with polarized 590 nm light. Note that, despite the red dye, vessels are visible; it is even possible to note where they descend into the parenchyma. Color-coded and numbered targets are shown in this view as well.

**Figure 12:**
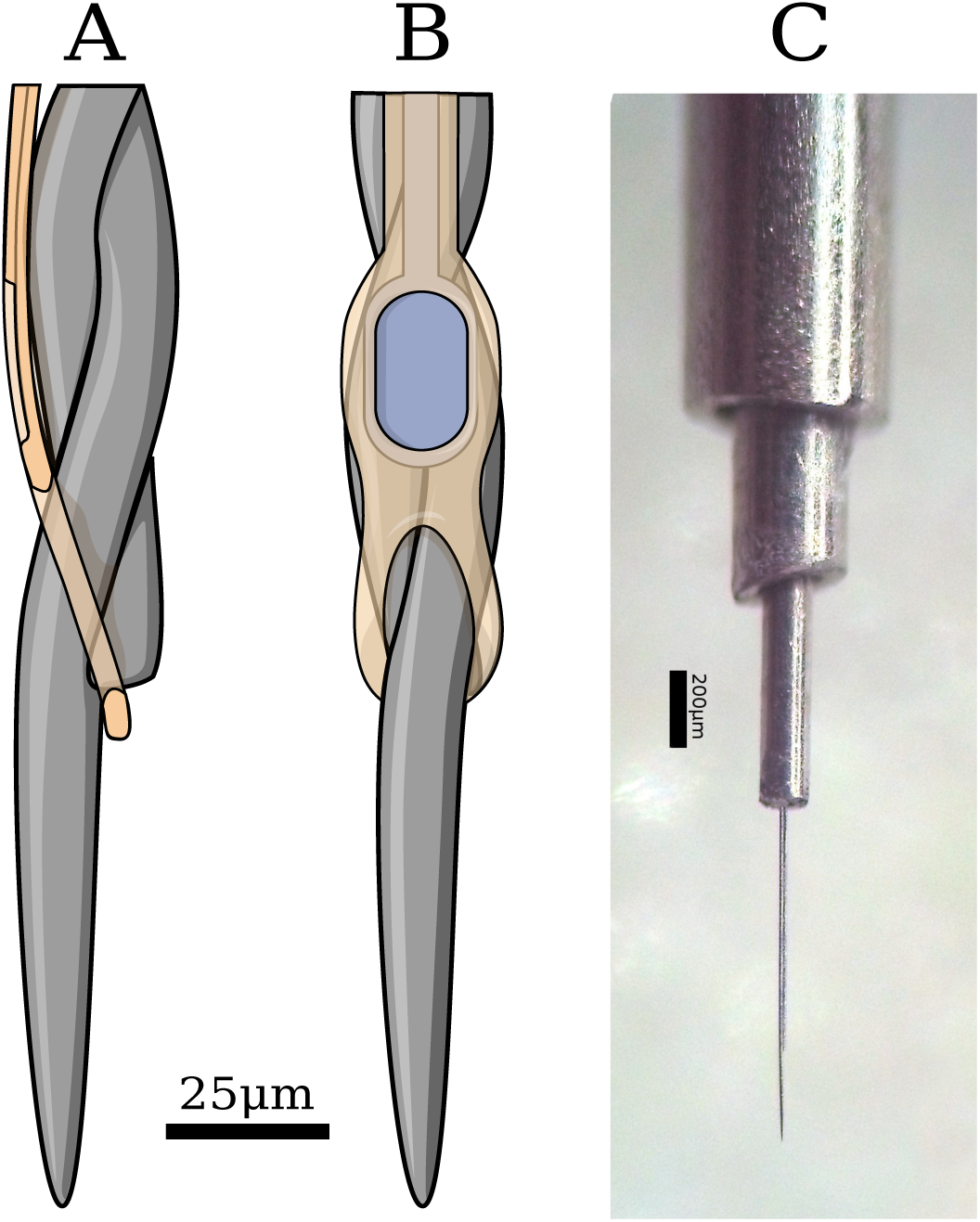
The inserter needle. Gray is tungsten-rhenium, amber is polyimide, blue is platinum. A. Side view, showing the position of the electrode loop on the needle step. B. Front view. Note the helix angle of the twisted wires is exaggerated for clarity. C. Micrograph of a fabricated needle in the telescoping needle cannula cartridge system.

## 8. Supplementary Materials

### 8.1. Software

As shown in Figure 8.1, the control software consists of 5 independent programs which communicate via Unix sockets.

’Video’ captures frames from the 4 cameras: 2 needle-electrode targeting, 90° and 45°, one targeting, and one oblique, and displays them on a monitor, with a UI for adding, moving, removing, zooming on, and sorting insertion targets as well as UI for selecting computer-vision templates for the needle and electrodes. ‘Register’ in turn uses these templates for locating the image positions of the respective elements; frames for this location are transmitted via shared memory. An example image of the cortical surface from ‘video’ is shown in figure 11.

’Register’ works by first bandpassing the image to minimize the effects of local luminance changes, then by iterative stochastic minimization of the computed difference between rotated and translated templates and video frames. As this is done using CUDA-accelerated texture access, over one million evaluations can be computed per second (40 Gpx/sec), objects can be located robustly within a few frames.

’Control’ uses the located positions to drive the needle into the electrode loop. Control in turn features a custom interpreted domain-specific language (DSL), which allows for rapid experimentation with e.g. different motor primitives for controlling the robot. Importantly, this DSL features persistent state between parsing, so that the DSL can be continually changed and refined during the course of a surgery. As this makes the experimenter more liable to err and crash the robot into itself or other things, the DSL features verification: before each segment of a motor primitive is executed, it is simulated, and checked against parametric limits. These limits are, in turn, directly written in the DSL, hence can be on primary variables (e.g. the position of each of the robots axes) or computed variables (e.g. variables modeling the mechanical chaining of the primary and secondary axes, or the computed target locations). This feature has proven invaluable for rapid and safe iteration, thereby maximizing the value of an animals life.

’Banger’ takes care of manually driving each of the 14 axes of servo and stepper motors, and 4 boolean lines. It receives per-axis absolute movement commands, and converts them into a realtime series of pulse / direction signals with constant-acceleration profiles. It is also capable of generating movements with non-zero velocity control points, which is instrumental in making high-speed pause-free movements during the control cycle. ‘Banger’ is run as a real-time blocking process during pulse generation, though we have found that a real-time kernel is not required for reliable timing.

Finally, ‘bbpid’ controls the ballistic retraction mechanism. During the course of insertion, bbpid rapidly switches between torque control (the spring loading phase), velocity (when slowing down the needle after retraction from the brain), and PID modes (holding position at top and bottom of stroke).

### 8.2. Inserter Needle

The needle is as critical to the system as any other element of the system; it needs to penetrate the brain, engage the electrode loop, and disengage during retraction. It is fabricated in a process not dissimilar to that used for tetrodes; however, instead of melted insulation, the four wires are joined with a Cu-Ir alloy, and the wires stop at different lengths.

First, one length of 12.5 *µm* W-26%Re wire is wound in a custom jig forming a ‘W’. One end of the W, which consists of two loops, is free to turn, and this allows all 4 wires to be twisted into a helix. The other end has the two fixed ends, held in brass clamps, and a loop that’s free to turn; this allows the 2 wires to form a continuation of the 4-helix. See Figure 16 for an illustration of the process. Both loops are held under tension via small springs, and the loops are rotated so that the helix angle of the 4 wires and 2 wires is approximately equal. Thus, two wires (the fixed ends) break off from the helix 1–10 mm before the end loop.

This jig has the added feature that the two ends are electrically isolated. The needle-winding jig is then installed in a chamber that is brought down to a vacuum of *<* 10 mTorr, before being filled with a shielding gas of 20% *H*_2_ in *Ar* at a pressure of 500 – 600 mTorr. See Figure 13 for a rendered cross-section through the brazing chamber, and Figure 14 for a photo of the exterior. Then, 100 mA of current is passed through the needle wires until they are *>* 1500 °C, at which point the oxides on the surface of the wire are either reduced or sublimate, leaving a clean metal surface. This temperature is adjusted to recrystalize the wire, as needed to create the step features. Following this, a mass of copper (group 11 elements are immiscible with tungsten, and copper is cheapest and has the highest boiling point) is melted in a boron-nitride crucible (carbon contaminates the melt and encourages the formation of tungsten carbide), and is raised to meet the wire helix. Copper has a very high contact angle with boron-nitride, which means that the meniscus of molten Cu can significantly exceed the lip of the crucible, critical for brazing. The needle-winding jig is then smoothly moved laterally several times, such that the molten braze metal joins the multiple wires into a single part. See Figure 15 for a photo of the interior of the brazing chamber and melt. We have experimented with dissolving iridium in the Cu melt to stiffen the braze alloy via solid solution strengthening, but have not had the time to demonstrate an effect, other than dramatically changing the surface morphology of the melt after gradual cooling (the surface of the braze on the needle is grassy due to rapid cooling and precipitation).

**Figure 13:**
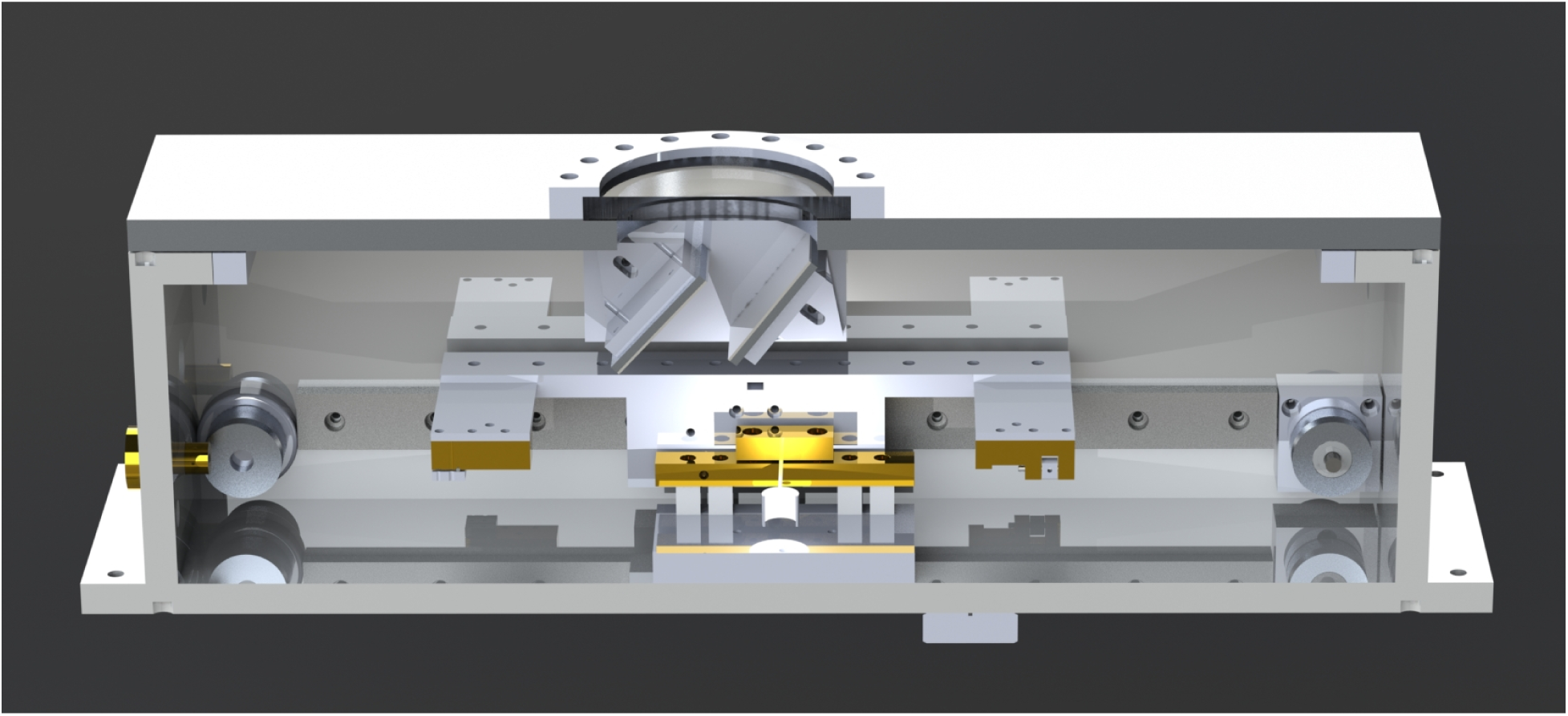
Render of a cross-section through the needle brazing machine. Jig A is installed on a linear slide, B, which moves laterally under control of a pulley system C attached to the end of vacuum feed-throughs. The melt, D, is held in a tungsten heater basket, in turn mounted to heavy current busses E (in turn fed by vacuum feed-throughs) which are mounted on a base which can move vertically, F. This permits the user to dip the wire into the melt under visual control through the viewing maze H, which prevents copper from condensing on the viewing window I in the lid.

**Figure 14:**
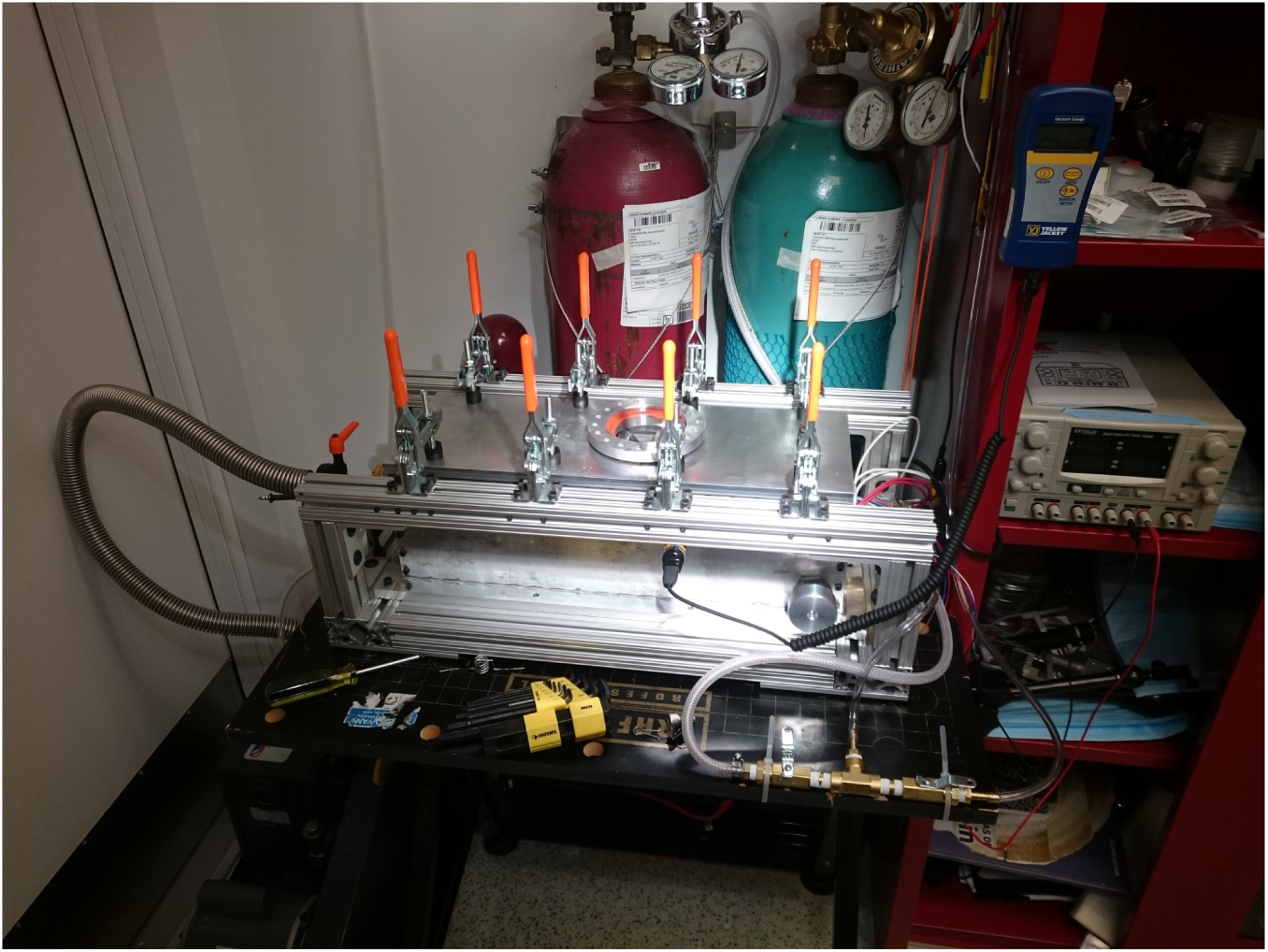
Photograph of the assembled, working needle brazing machine.

**Figure 15:**
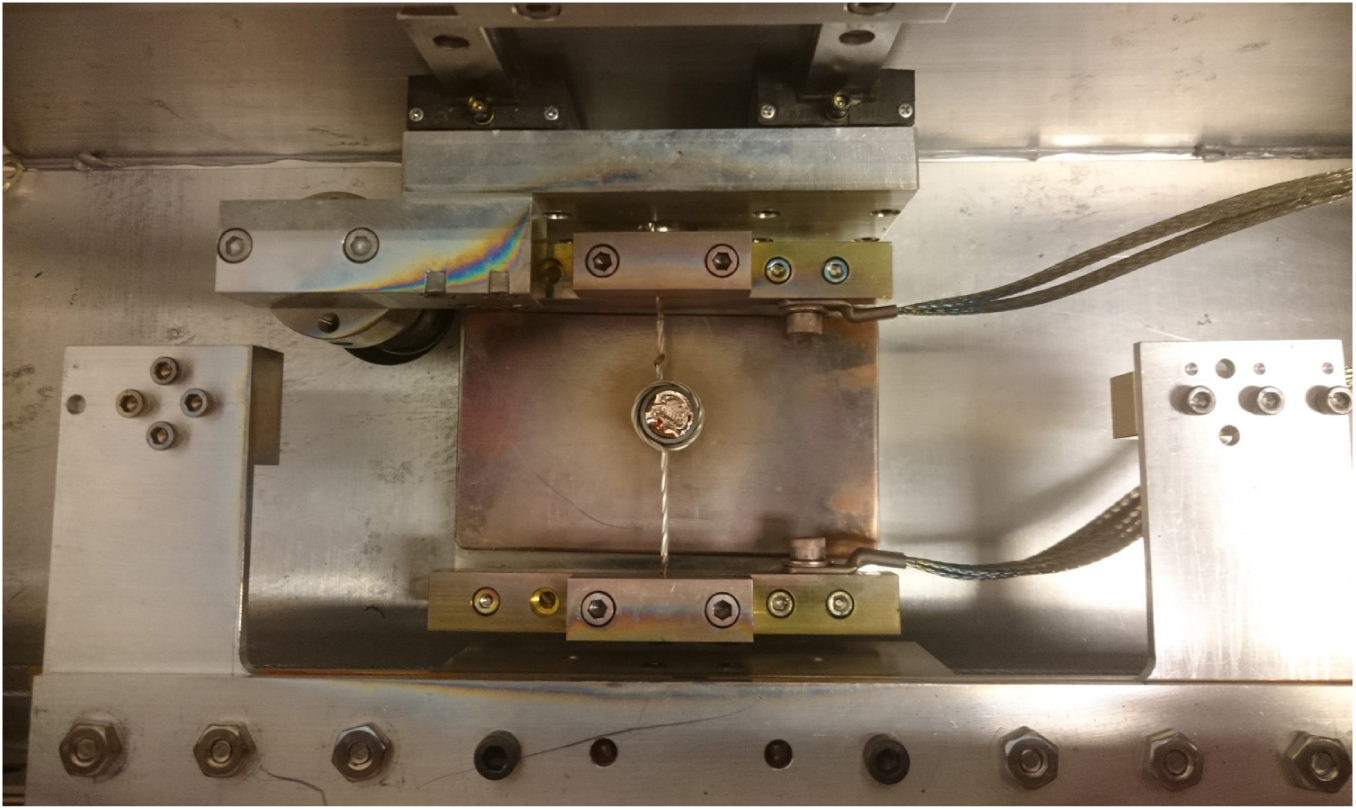
Photograph of the interior of the brazing machine, with the copper melt and needle jig installed.

The nascent needle is then removed from the molten braze material, the heating elements to the crucible are disengaged, and the vacuum chamber is allowed to cool prior being flushed with argon. The chamber is then opened, and the needle-winding jig is removed. Then, three of the four wires that exit the helix are removed by fatiguing the wire at the point where it leaves the helix. Two of these wires exit the helix 1-10 mm from the last, and provide strength to prevent buckling within the telescoping cannula. As mentioned above, the annealing / reducing temperature is critical here – too low, and the wire retains its as-drawn ductile nature, which will lead to a barb, not a step; too high, and the whole needle is brittle, and likely to fracture during assembly and use. As-drawn, the wire has very elongated crystalline domains, which leads to ductility and high tensile strength of the material; these domains become more regular, and orient perpendicular to the direction of applied stress (tension due to springs in the needle-winding jig) during annealing and recrystalization. The addition of rhenium to the tungsten wire provides two advantages at this point: it raises the recrystalization temperature, so that the window between surface-oxide reduction and recrystalization is larger, and it increases the strength and modulus of the resulting needle.

Even with careful control of both annealing and brazing profiles, yield on the final step – that which engages the electrode – is low, primarily because with proper annealing / recrystalization, the shorter wire tends to break 10 *µm* from where the braze fillet ends. This leads to a ‘fork’ which the thin electrodes can be stuck in during insertion. To remedy this, a very fine sandpaper (e.g. 10 *µm* grit) is delicately run along the needle, with 3 wires removed, but one wire still attached to the needle-winding jig, until elements of the step are eroded to the point that no fork can be observed, and the needle is as drawn in Figure 12 (though with a much lower helix angle).

**Figure 16:**
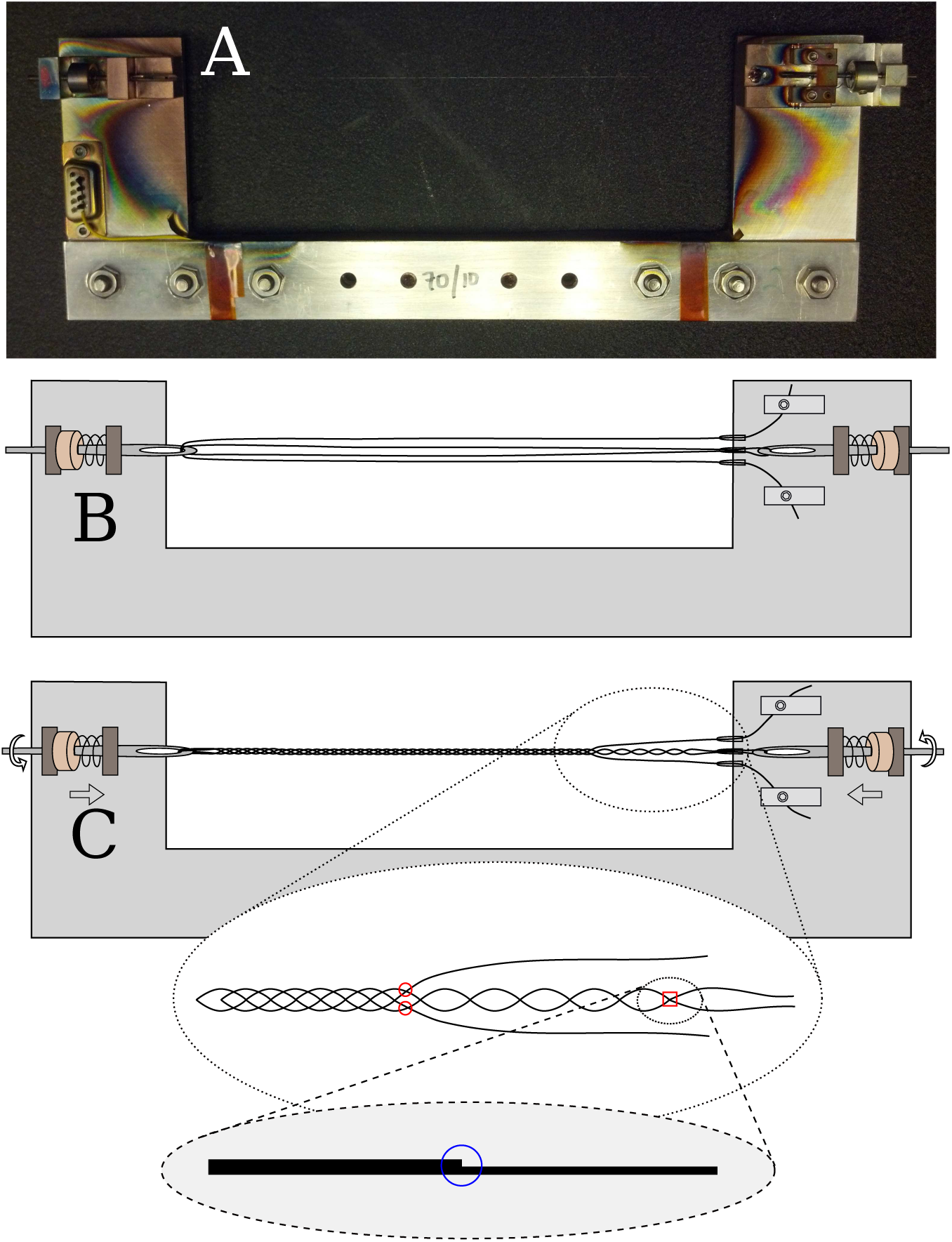
A. Photo of a needle brazing jig, with wire visible, prior brazing. Left is connector for supplying current to heat the needle for surface oxide reduction and recrystalization. Visible copper coating is from melt evaporation. B. Schematic of the jig configuration prior wire spinning, showing one length of wire pulled into a ‘W’ shape through the loops and eyelets. **C**. Schematic of the jig prior brazing, with details of where the first two wires break off, and a detail where the third wire separates to form a step.

At this point, the needle is loaded into a quadruple-telescoping cartridge, the far end is attached to the upper part of this cartridge, and this device is installed in another machine for inspecting and etching the needle, shown in Figure 17. This machine consists of a microscope and two etchant wells. The first is 0.2 M *FeCl*_3_, for removing Cu from the surface of the needle; the microscope is used to dip the needle into the solution just to the point of the shoulder / final break. The second is 0.2 M *NaOH*, which is used with 1-6V AC to electrolytically etch the longest part of the needle to a fine point. This machine also affords assessment of needle motion, which can be impeded if any dust or dirt entered the cannula during assembly.

**Figure 17:**
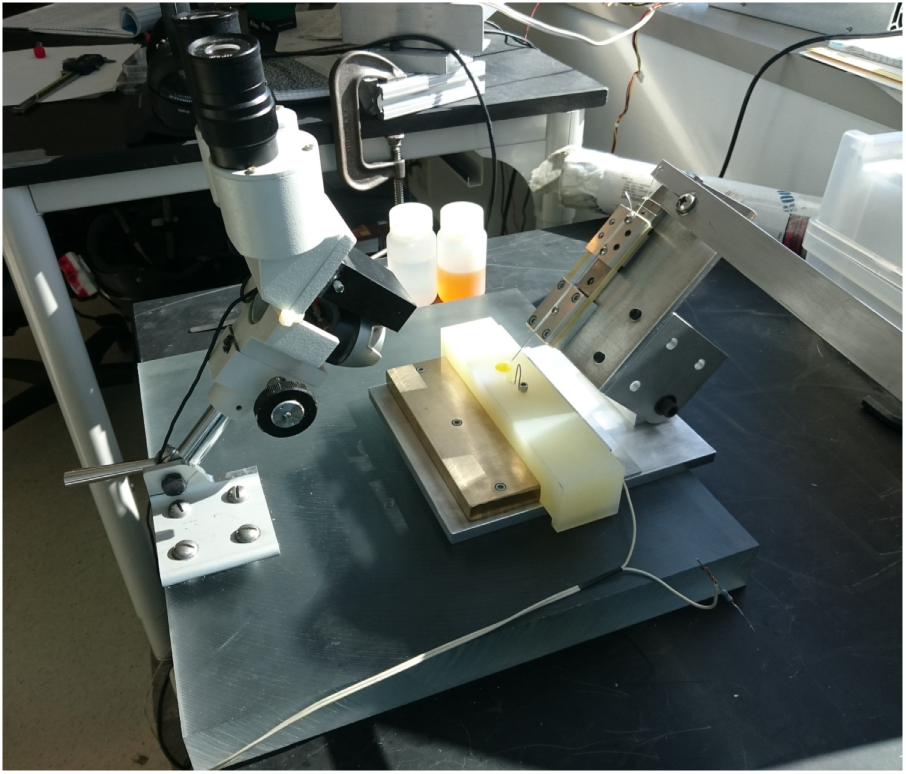
Photograph of the needle etching and inspection station.

### 8.3. Ballistic Retraction

The ballistic retraction mechanism serves to allow slow, precisely controlled insertion, with rapid high-acceleration retraction. As shown in Figure 18, this is accomplished with a slug + cylinder +nail-head system. The mechanism also permits the needle to be freely rotated, necessary as the needle step needs to be oriented away from the cartridge knife-edge. (In past robot revisions, needle rotation was used to disengage the electrode from the needle; this is no longer needed with ballistic retraction.)

**Figure 18:**
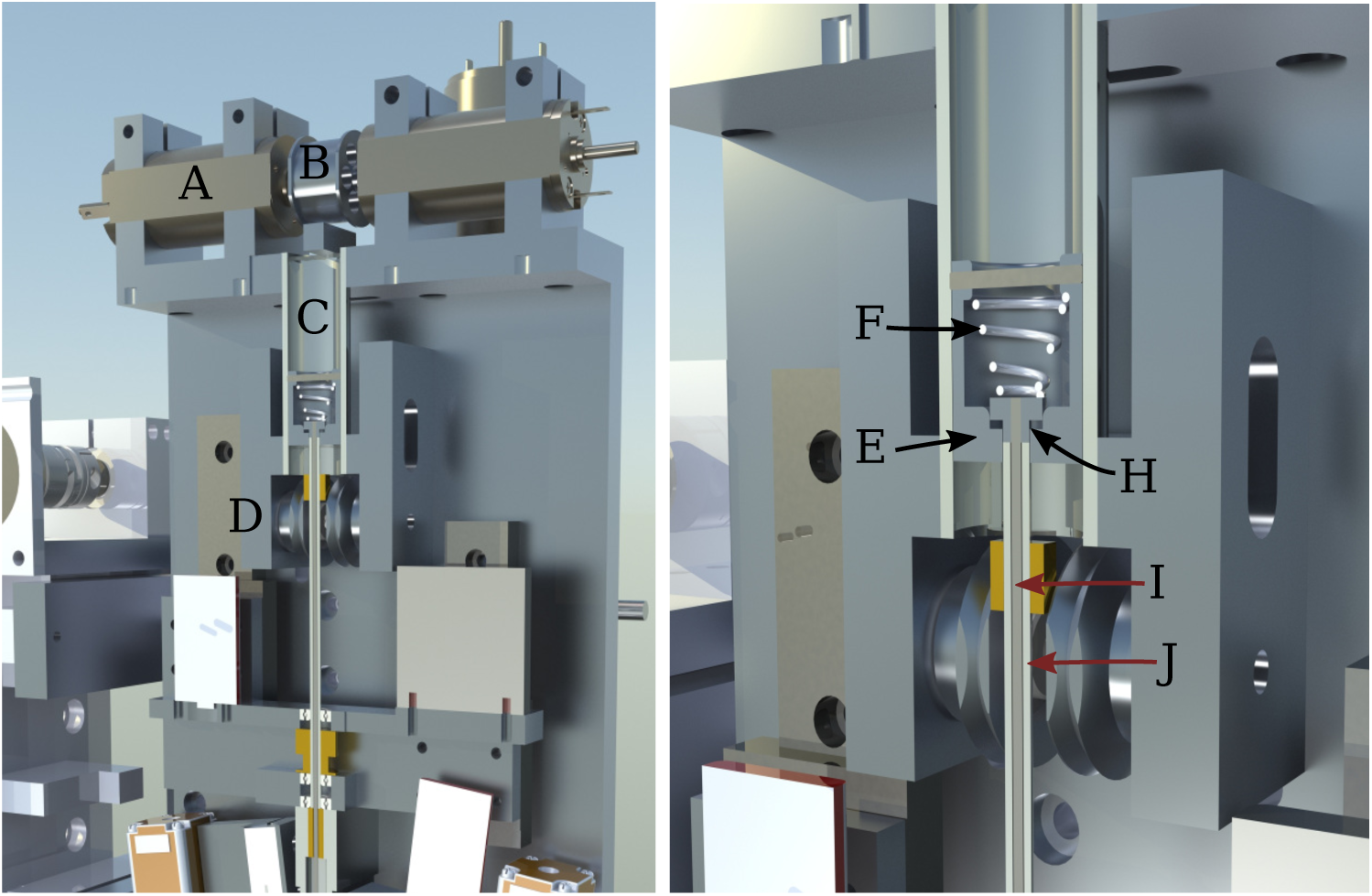
Overview render of the ballistic retraction mechanism. **A**. Two high-speed low-inertia Maxon motors drive a small pulley **B**. The pulley in turn drives a hollow slug E up and down inside a cylinder **C**.. Kevlar cables for actuating the slug are omitted in this rendering, but go around adjustable pulleys **D**. During the retraction phase, the motors pulse downward, compressing the spring **F**. Then the motors reverse, driving the slug E into the nail-head H, which is silver-nickel brazed onto the 17-4PH needle actuation rod **I**. This rod rides in a notched tube J, which allows precise control of needle insertion depth. After impact the slug is slowed to a stop at the upper position in the cylinder. During the subsequent insertion phase, the high-speed motors drive the slug lightly down, so that the insertion force is supplied by the spring and kevlar cables.

## Footnotes

‡ An upward-facing camera with 25 *µm* tungsten reticle wires is used to determine this calibration. First, the reticle is located in the targeting camera, and then the inserter head is moved over the upward facing camera, and the inner diameter of the needle cartridge is aligned to the reticle, thereby transferring the optical coordinate frame to the mechanical insertion head. It is also possible to transfer the coordinate frame from camera to robot by locating an electrode loop in the camera; when the needle is through the loop, robot position is known in both coordinate frames.

